# A mechanistic-statistical approach to infer dispersal and demography from invasion dynamics, applied to a plant pathogen

**DOI:** 10.1101/2023.03.21.533642

**Authors:** Méline Saubin, Jérome Coville, Constance Xhaard, Pascal Frey, Samuel Soubeyrand, Fabien Halkett, Frédéric Fabre

**Affiliations:** Université de Lorraine, INRAE, IAM, F-54000 Nancy, France; INRAE, BioSP, 84914 Avignon, France; Universitéde Lorraine, INSERM CIC-P 1433, CHRU de Nancy, INSERM U1116, Nancy, France; INRAE, Bordeaux Sciences Agro, SAVE, F-33882 Villenave d’Ornon, France

**Author notes:** **Corresponding authors:** Méline Saubin, Fabien Halkett, Frédéric Fabre. Populationsgenetik, Technische Universität München, Liesel-Beckmann-Str. 2, 85354 Freising, Germany (Méline Saubin). These authors co-directed this work (Fabien Halkett, Frédéric Fabre).

**Keywords:** 1-D colonisation, dispersal kernel, long-distance dispersal, multiple data types, population dynamic, spatio-temporal model

## Abstract

Dispersal, and in particular the frequency of long-distance dispersal (LDD) events, has strong implications for population dynamics with possibly the acceleration of the colonisation front, and for evolution with possibly the conservation of genetic diversity along the colonised domain. However, accurately inferring LDD is challenging as it requires both large-scale data and a methodology that encompasses the redistribution of individuals in time and space. Here, we propose a mechanistic-statistical framework to estimate dispersal from one-dimensional invasions. The mechanistic model takes into account population growth and grasps the diversity in dispersal processes by using either diffusion, leading to a reaction-diffusion (R.D.) formalism, or kernels, leading to an integro-differential (I.D.) formalism. The latter considers different dispersal kernels (*e*.*g*. Gaussian, Exponential, and Exponential-power) differing in their frequency of LDD events. The statistical model relies on dedicated observation laws that describe two types of samples, clumped or not. As such, we take into account the variability in both habitat suitability and occupancy perception. We first check the identifiability of the parameters and the confidence in the selection of the dispersal process. We observed good identifiability for all parameters (correlation coefficient *>* 0.9 between true and fitted values). The dispersal process that is the most confidently identified is Exponential-Power (*i*.*e*. fat-tailed) kernel. We then applied our framework to data describing an annual invasion of the poplar rust disease along the Durance River valley over nearly 200 km. This spatio-temporal survey consisted of 12 study sites examined at seven time points. We confidently estimated that the dispersal of poplar rust is best described by an Exponential-power kernel with a mean dispersal distance of 1.94 km and an exponent parameter of 0.24 characterising a fat-tailed kernel with frequent LDD events. By considering the whole range of possible dispersal processes our method forms a robust inference framework. It can be employed for a variety of organisms, provided they are monitored in time and space along a one-dimension invasion.

## 1 Introduction

Dispersal is key in ecology and evolutionary biology (Clobert et al., 2004). From an applied point of view, the knowledge of dispersal is of prime interest for designing ecological-based management strategies in a wide diversity of contexts ranging from the conservation of endangered species (*e*.*g*., Macdonald and Johnson, 2001) to the mitigation of emerging epidemics (Dybiec et al., 2009; Fabre et al., 2021). From a theoretical point of view, the pattern and strength of dispersal sharply impact eco-evolutionary dynamics (*i*.*e*. the reciprocal interactions between ecological and evolutionary processes) (Miller et al., 2020). The features of dispersal have many implications for population dynamics (*e*.*g*. speed of invasion, metapopulation turnover; Soubeyrand et al., 2015; Kot et al., 1996), genetic structure (*e*.*g*. gene diversity, population differentiation; Edmonds et al., 2004; Fayard et al., 2009; Petit, 2011) and local adaptation (Gandon and Michalakis, 2002; Hallatschek and Fisher, 2014). Mathematically, the movement of dispersers (individuals, spores or propagules for example) can be described by a so-called location dispersal kernel (Nathan et al., 2012) that represents the statistical distribution of the locations of the propagules of interest after dispersal from a source point. Since the pioneer works of Mollison (1977), much more attention has been paid to the “fatness” of the tail of the dispersal kernel (Klein et al., 2006). Short-tailed kernels (also referred to as “thin-tailed”) generate an invasion front of constant velocity, whereas long-tailed kernels (also referred to as “fat-tailed”) can cause an accelerating front of colonisation (Ferrandino, 1993; Kot et al., 1996; Clark et al., 2001; Mundt et al., 2009; Hallatschek and Fisher, 2014). Long-tailed kernels, characterised by more frequent long-distance dispersal (LDD) events than an exponential kernel that shares the same mean dispersal distance, can also cause a reshuffling of alleles along the colonisation gradient, which prevents the erosion of genetic diversity (Nichols and Hewitt, 1994; Petit, 2004; Fayard et al., 2009) or leads to patchy population structures (Ibrahim et al., 1996; Bialozyt et al., 2006).

Despite being a major issue in biology, properly characterising the dispersal kernels is a challenging task for many species, especially when dispersing individuals are numerous, small (and thus difficult to track) and move far away (Nathan, 2001). In that quest, mechanistic-statistical models enable a proper inference of dispersal using spatio-temporal datasets (Wikle, 2003a; Soubeyrand et al., 2009a; Roques et al., 2011; Soubeyrand and Roques, 2014; Hefley et al., 2017; Nembot Fomba et al., 2021) while allowing for the parsimonious representation of both growth and dispersal processes in heterogenous environments (Papaїx et al., 2022). They require detailed knowledge of the biology of the species of interest to properly model the invasion process. They combine a mechanistic model describing the invasion process and a probabilistic model describing the observation process while enabling a proper inference using spatio-temporal data. Classically, the dynamics of large populations are well described by deterministic differential equations. Invasions have often been modelled through reaction-diffusion equations (Murray, 2002; Okubo and Levin, 2002; Shigesada and Kawasaki, 1997). In this setting, individuals are assumed to move randomly following trajectories modelled using a Brownian motion or a more general stochastic diffusion process. Despite their long standing history, the incorporation of reaction-diffusion equations into mechanistic-statistical approaches to estimate parameters of interest from spatio-temporal data essentially dates back to the early 2000s (*e*.*g*. Wikle, 2003a; Soubeyrand and Roques, 2014; Louvrier et al., 2020; Nembot Fomba et al., 2021). By contrast to reaction-diffusion equations, integro-differential equations encode trajectories modelled by jump diffusion processes and rely on dispersal kernels, individuals being redistributed according to the considered kernel (Fife, 1996; Hutson et al., 2003; Kolmogorov et al., 1937). This approach allows to consider a large variety of dispersal functions, typically with either a short or a long tail (*i*.*e*. putative LDD events). As such it is more likely to model accurately the true organism’s dispersal process. In the presence of long-distance dispersal, the biological interpretation of the estimated diffusion parameters with an R.D. equation would be misleading. However, integro-differential equations are numerically more demanding to simulate than reaction-diffusion equations. As far as we know, integro-differential equations have rarely been embedded into mechanistic-statistical approaches to infer dispersal processes in ecology (but see Szymańska et al., 2021 for a recently proposed application of a non-local model to cell proliferation).

Data acquisition is another challenge faced by biologists in the field, all the more that data confined to relatively small spatial scales can blur the precise estimates of the shape of the kernel’s tail (Ferrandino, 1996; Kuparinen et al., 2007; Rieux et al., 2014). To gather as much information as possible, it is mandatory to collect data over a wide range of putative population sizes (from absence to near saturation) along the region of interest. Sharing the sampling effort between raw and refined samples to browse through the propagation front may improve the inference of spatial ecological processes (Gotway and Young, 2002). This way of sampling is all the more interesting as the probabilistic model describing the observation process in the mechanistic-statistical approach can handle such multiple datasets (Wikle, 2003b). However, inference based on multi-type data remains a challenging statistical issue as the observation variables describing each data type follow different distribution laws (Chagneau et al., 2011) and can be correlated or, more generally, dependent because they are governed by the same underlying dynamics (Bourgeois et al., 2012; Georgescu et al., 2014; Soubeyrand et al., 2018). This requires a careful definition of the conditional links between the observed variables and the model parameters (the so-called observation laws) in order to identify and examine complementarity and possible redundancy between data types.

In this article, we aim to provide a sound and unified inferential framework to estimate dispersal from ecological invasion data using both reaction-diffusion and integro-differential equations. We first define the two classes of mechanistic invasion models, establish the observation laws corresponding to raw and refined samplings, and propose a maximum-likelihood method to estimate their parameters within the same inferential framework. Then, to confirm that each model parameter can indeed be efficiently estimated given the amount of data at hand (see Soubeyrand and Roques, 2014), we perform a simulation study to check model parameters’ identifiability given the sampling design. We also aim to assess the confidence level in the choice of the dispersal function as derived by model selection. Last, the inferential framework is applied to original ecological data describing the annual invasion of a tree pathogen (*Melampsora larici-populina*, a fungal species responsible for the poplar rust disease) along the riparian stands of wild poplars bordering the Durance River valley in the French Alps (Xhaard et al., 2012).

## 2 Modelling one-dimensional invasion and observation processes

### 2.1 A class of deterministic and mechanistic invasion models

We model the dynamics of a population density *u*(*t, x*) at any time *t* and point *x* during an invasion using two types of spatially heterogeneous deterministic models allowing to represent a wide range of dispersal processes. Specifically, we considered a reaction-diffusion model (R.D.) and an integro-differential model (I.D.):

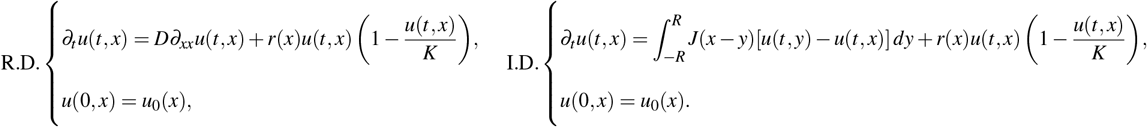

where *t* varies in [0, *T*] (*i*.*e*. the study period) and *x* varies in [*−R, R*] (*i*.*e*. the study domain). Both equations exhibit the same structure composed of a diffusion/dispersal component and a reaction component. The reaction component, 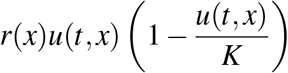 in both equations, is parameterised by a spatial growth rate *r*(*x*) that takes into account macro-scale variations of the factors regulating the population density and *K* the carrying capacity of the environment. It models population growth. The diffusion/dispersal component models population movements either by a diffusion process (*D∂*_*xx*_*u* in R.D.) parameterised by the diffusion coefficient *D* or by a dispersal kernel (*J* in I.D.). To cover a large spectrum of possible dispersal processes, we use the following parametric form for the kernel *J*:

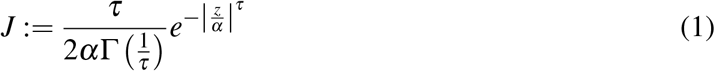

with mean dispersal distance 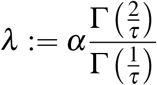 . Varying the value of *τ* leads to the kernels classically used in dispersal studies. Specifically, *J* can be a Gaussian kernel 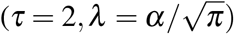, an exponen tial kernel (*τ* = 1, *λ* = *α*) or a fat-tail kernel 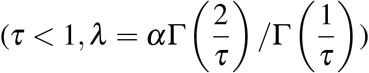. Explicit formulas for the solution *u*(*t, x*) of these reaction-diffusion/dispersal equations being out of reach, we compute a numerical approximation *u*_num_ of *u*, which serves as a surrogate for the real solution. Details of the numerical scheme used to compute *u*_num_ can be found in Appendix S1.

### 2.2 A conditional stochastic model to handle micro-scale fluctuations

Among the factors driving population dynamics, some are structured at large spatial scales (macro-scale) and others at local scales (micro-scale). It is worth considering both scales when studying biological invasions. In the model just introduced, the term *r*(*x*) describes factors impacting population growth rate at the macro-scale along the whole spatial domain considered. Accordingly, the function *u*(*t, x*) is a mean-field approximation of the true population density at macro-scale. Furthermore, the population density can fluctuate due to micro-scale variations of other factors regulating population densities locally (*e*.*g*. because of variations in the micro-climate and the host susceptibility). Such local fluctuations are accounted for by a conditional probability distribution on *u*(*t, x*), the macro-scale population density, which depends on the (unobserved) suitability of the habitat unit as follow. Consider a habitat unit *i* whose centroid is located at *x*_*i*_, and suppose that the habitat unit is small enough to reasonably assume that *u*(*t, x*) = *u*(*t, x*_*i*_) for every location *x* in the habitat unit. Let *N*_*i*_(*t*) denote the number of individuals in *i* at time *t*. The conditional distribution of *N*_*i*_(*t*) is modelled by a Poisson distribution:

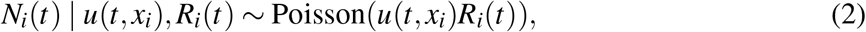

where *R*_*i*_(*t*) is the intrinsic propensity of the habitat unit *i* to be occupied by individuals of the population at time *t*. Thereafter, *R*_*i*_(*t*) is called habitat suitability and takes into account factors like the exposure and the favorability of habitat unit *i*. The suitability of habitat unit *i* is a random effect (unobserved variable) and is assumed to follow a Gamma distribution with shape parameter *σ*^*−*2^ and scale parameter *σ* ^2^:

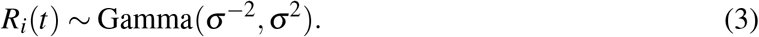

This parametrisation implies that the mean and variance of *R*_*i*_(*t*) are 1 and *σ* ^2^, respectively; that the conditional mean and variance of *N*_*i*_(*t*) given *u*(*t, x*_*i*_) are *u*(*t, x*_*i*_) and *u*(*t, x*_*i*_) + *u*(*t, x*_*i*_)^2^*σ* ^2^, respectively; and that its conditional distribution is:

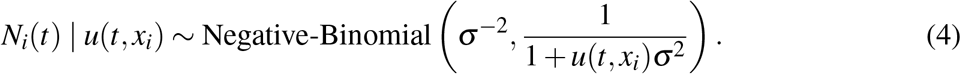

### 2.3 Multi-type sampling and models for the observation processes

During an invasion, the population density may range from zero (beyond the front) to the maximum carrying capacity of the habitat. To optimise the sampling effort, it may be relevant to carry out different sampling procedures depending on the population density at the sampling sites. In this article, we consider a two-stage sampling made of one raw sampling, which is systematic and one optional refined sampling adapted to our case study, the downstream spread of a fungal pathogen along a river (Figure 1). We consider that the habitat unit is a leaf. The fungal population is monitored in sampling sites *s* ∈ {1,…, *S*} and at sampling times *t* ∈ {*t*_1_, …, *t*_*K*_}. Sampling sites are assumed to be small with respect to the study region, and the duration for collecting one sample is assumed to be short with respect to the study period. Thus, the (macro-scale) density of the population at sampling time *t* in sampling site *s* is constant and equal to *u*(*t, z*_*s*_) where *z*_*s*_ is the centroid of the sampling site *s*. Any sampling site *s* is assumed to contain a large number of leaves which are, as a consequence of the assumptions made above, all associated with the same population density function: *u*(*t, x*_*i*_) = *u*(*t, z*_*s*_) for all leaves *i* within sampling site *s*. Each observed tree and twig are assumed to be observed only once during the sampling period. Therefore, habitat suitabilities *R*_*i*_(*t*) are considered independent in time.

**Figure 1.**
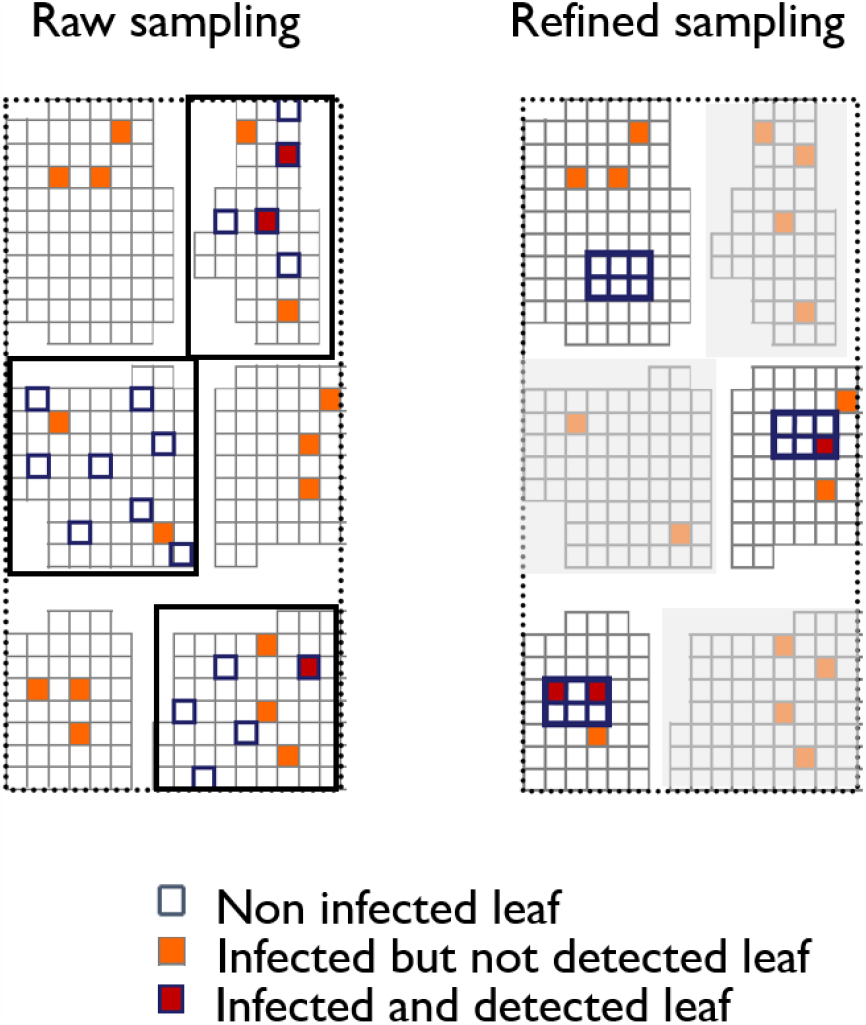
Two-stage sampling on a sampling site, with one systematic raw sampling (on the left) and one optional refined sampling (on the right). Each square represent a leaf, which can be non infected, infected but not detected, or infected and detected. Each group of spatially grouped leaves represent a tree. Each tree already observed during the raw sampling are not available (and thus represented in grey) for the refined sampling, where connected leaves in twigs are observed.

The raw sampling is focused on trees, considered as a group of independent leaves regarding their suitabilities. This assumption can be made if the leaves observed on the same tree are sufficiently far from each other and represent a large variety of environmental conditions, and therefore habitat suitabilities (for example, leaves observed all around a tree will not have the same sun exposition, nor the same humidity depending on their height and their relative positions to the trunk). In each sampling site *s* and at each sampling time *t*, a number *B*_*st*_ of trees are monitored for the presence of infection. We count the number of infected trees *Y*_*st*_ among the total number *B*_*st*_ of observed trees. In the simulations and the case study tackled below, the random variables *Y*_*st*_ given *u*(*t, x*_*s*_) are independent and distributed under the conditional Binomial distribution 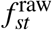 described in Appendix S2.2. Its success probability depends on the variabilities of (i) the biological process through the variance parameter *σ*^2^ of habitat suitabilities, and (ii) the observation process through a parameter *γ*. This parameter describes how the probabilities of leaf infection perceived by the person in charge of the sampling differ between trees from true probabilities (as informed by the mechanistic model). Such differences may be due, for example, to the specific configuration of the canopy of each tree or to particular lighting conditions.

The refined sampling is focused on twigs, considered as a group of connected leaves. Nearby leaves often encounter the same environmental conditions and, therefore, are characterised by similar habitat suitabilities represented by *R*_*i*_(*t*); see Equations (2–3). This spatial dependence was taken into account by assuming that the leaves of the same twig (considered as a small group of spatially connected leaves) share the same leaf suitability. Accordingly, suitabilities are considered as shared random effects. The refined sampling is performed depending on disease prevalence and available time. In site *s* at time *t, G*_*st*_ twigs are collected. For each twig *g*, the total number of leaves *M*_*stg*_ and the number of infected leaves *Y*_*stg*_ are counted. In the simulations and the case study tackled below, the random variables *Y*_*stg*_ given *u*(*t, x*_*s*_) are independent and distributed under conditional probability distributions denoted by 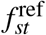 described in Appendix S2.3. The distribution 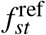 is a new mixture distribution (called Gamma-Binomial distribution) obtained using Equations (2–3) and taking into account the spatial dependence and the variance parameter of unobserved suitabilities (see Appendix S2.3).

This sampling scheme and its vocabulary (leaves, twigs and trees) are specifically adapted to our case study for the sake of clarity. However, a wide variety of multi-type sampling strategies can be defined and implemented in the model, as long as it fits a two-stage sampling as presented in Figure 1.

### 2.4 Coupling the mechanistic and observation models

The submodels of the population dynamics and the observation processes described above can be coupled to obtain a mechanistic-statistical model (also called physical-statistical model; Berliner, 2003; Soubeyrand et al., 2009b) representing the data and depending on dynamical parameters, namely the growth and dispersal parameters. The likelihood of this mechanistic-statistical model can be written:

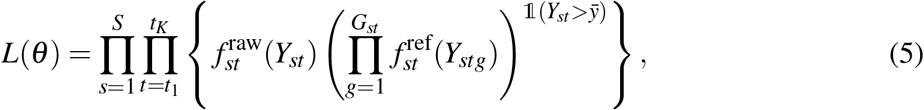

where 1(*·*) denotes the indicator function and expressions of 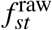 and 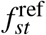adapted to the case study tackled below are given by Equations (S14) and (S18) in Appendix S2. The power 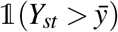 equals to 1 if 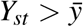 and 0 otherwise, implies that the product 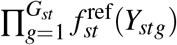 only appears if the refined sampling is carried out in site *s*. Moreover, such a product expression for the likelihood is achieved by assuming that leaves in the raw sampling and those in the refined sampling are not sampled from the same trees. If this does not hold, then an asymptotic assumption like the one in Appendix S2.2 can be made to obtain Equation (5), or the dependence of the unobserved suitabilities must be taken into account and another likelihood expression must be derived.

## 3. Parameter estimation and model selection

We performed simulations to check the practical identifiability of several scenarios of biological invasions. Invasion scenarios represent a wide range of possible states of nature regarding the dispersal process, the environmental heterogeneity at macro-scale, and the intensity of local fluctuations at micro-scale. Even though the simulations are designed to cope with the structure of our real data set (Appendix S4), the results enable some generic insights to be gained. Specifically, we considered six sampling dates evenly distributed in time and 12 samplings sites evenly distributed within the 1D spatial domain. For each pair *(date, site)*, we simulated the raw sampling of 100 trees and the refined sampling of 20 twigs. For the fifth sampling date, the raw sampling was densified with 45 sampling sites instead of 12.

The simulation study explored four hypotheses for the dispersal process: three I.D. hypotheses with kernels *J*_Exp_, *J*_Gauss_ and *J*_ExpP_ and the R.D. hypothesis. Hypotheses *J*_Exp_ and *J*_Gauss_ state that individuals dispersed according to Exponential and Gaussian kernels, respectively, with parameter *θ*_*J*_ = (*λ*). Hypothesis *J*_ExpP_ states that individuals dispersed according to a fat-tail Exponential-power kernel with parameters *θ*_*J*_ = (*λ, τ*) and *τ <* 1. Finally, hypothesis R.D. states that individual dispersal is a diffusion process parameterised by *θ*_*J*_ = (*λ*). The parameter *λ* represents the mean distance travelled whatever the dispersal hypothesis considered. Moreover, macro-scale environmental heterogeneity was accounted for in the simulations by varying the intrinsic growth rate of the pathogen population (*r*) in space. Specifically, along the one-dimensional domain, we considered two values of *r*, namely a downstream value *r*_dw_ and an upstream value *r*_up_, parameterised by *θ*_*r*_ = (*r*_dw_, *ω*) such that 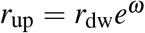 . Finally, micro-scale heterogeneity was accounted for in the simulations by varying the parameter of leaf suitability *σ* ^2^ and tree perception *γ*. Thereafter, *θ* = (*θ*_*r*_, *θ*_*J*_, *γ, σ* ^2^) denotes the vector of model parameters.

### 3.1 Accurate inference of model parameters

To assess the estimation method and check if real data that were collected are informative enough to efficiently estimate the parameters of the models (the so-called practical identifiability), we proceeded in three steps for each dispersal hypothesis: (i) a set of parameter values *θ* = (*θ*_*r*_, *θ*_*J*_, *γ, σ* ^2^) is randomly drawn from a distribution that encompasses a large diversity of realistic invasions, (ii) a data set with a structure similar to our real sampling is simulated given *θ* and (iii) *θ* is estimated using the maximum-likelihood method applied to the simulated data set. These steps were repeated *n* = 160 times. Details on the simulation procedure, the conditions used to generate realistic invasions, and on the estimation algorithm are provided in Appendix S4.1. Practical identifiability was tested by means of correlation coefficients between the true and estimated parameter values (see Table 1, Appendix S2: Figures S2, S3, S4, S5).

**Table 1:** Model practical identifiability. Numbers indicate the coefficient of correlation between the true and estimated parameter values for the four models corresponding to the four dispersal processes (*J*_Exp_, *J*_Gauss_, *J*_ExpP_ and R.D.) from 160 replicates. High correlation between true and estimated parameters indicates a good practical identifiability. The standard deviations of the coefficients of correlation, estimated with a bootstrapping method, are indicated in brackets. Correlation coefficients and standard deviations are given for natural scale for parameter *ω*, and logarithm scales for parameters *r*_dw_, *γ, λ, τ*, and *σ* ^2^.

| Parameter | Description | $J_{\text{Exp}}$ | $J_{\text{Gauss}}$ | $J_{\text{ExpP}}$ | R.D. |
| --- | --- | --- | --- | --- | --- |
| $r_{\text{dw}}$ | Growth rate downstream | 0.997 (0.001) | 0.997 (0.001) | 0.992 (0.004) | 0.991 (0.003) |
| $\omega$ | Growth rate modulator | 0.997 (0.001) | 0.992 (0.003) | 0.994 (0.002) | 0.995 (0.002) |
| $\lambda$ | Mean dispersal distance | 0.983 (0.007) | 0.993 (0.004) | 0.983 (0.006) | 0.941 (0.023) |
| $\tau$ | Kernel exponent | NA | NA | 0.927 (0.019) | NA |
| $\gamma$ | Tree perception | 0.969 (0.006) | 0.966 (0.006) | 0.966 (0.006) | 0.910 (0.018) |
| $\sigma^2$ | Variance in leaf suitability | 0.996 (0.001) | 0.997 (0.001) | 0.997 (0.001) | 0.994 (0.001) |

All the parameters defining the macro-scale mechanistic invasion model (*r*_*dw*_, *ω, λ*) display very good practical identifiability whatever the model, with correlation coefficients above 0.98 (except for mean dispersal distance *λ* under R.D., correlation coefficient of 0.94). In the case of the Exponential-power dispersal kernel, the additional parameter representing the tail of the distribution (*τ*) also displays a very good practical identifiability with a correlation coefficient of 0.93. The parameter defining the micro-scale fluctuations, *σ*^2^, leads to particularly high correlation coefficients (0.99 for all the models). The identifiability for the perception parameter *γ* related to the observation process is somewhat lower (from 0.91 to 0.97).

### 3.2 Confidence in the selection of the dispersal process

Numerical simulations were next designed to test whether model selection could disentangle the true dispersal process (*i*.*e*. the dispersal hypothesis used to simulate the data set) from alternative dispersal processes (Appendix S4.2). The model selection procedure is the most efficient for the dispersal hypotheses Exponential-power *J*_ExpP_, with 78% of correct kernel selection, respectively (Table 2). When the fat-tail Exponential-power kernel is not correctly identified, it is mostly mistaken with the Exponential one (for 17% of the simulations). In line with this, the probability of correctly selecting the kernel *J*_ExpP_ decreases when the parameter *τ* increases towards 1, the value for which the Exponential-power kernel coincides with the Exponential kernel (Figure 2). Importantly, when the Exponential-power kernel is correctly selected, we observe a large difference between its AIC and the AIC of the second best model (217.50 points on average). Conversely, when the invasion process is simulated under *J*_ExpP_, but another kernel is selected, we observe a very small AIC difference (0.76 point on average). The model selection is the least efficient for the Gaussian kernel *J*_Gauss_ (Table 2), with only 45% of correct model selection. Its correct identification is improved to 80% by densifying the sampling scheme (Appendix S4.5: Table S2). Finally, note that when the invasion process is simulated under model R.D. or *J*_Gauss_, a short-tail kernel is almost always selected and, thus, never confounded with the fat-tail kernel *J*_ExpP_.

**Table 2:** Efficiency of model selection using Akaike information criterion (AIC). The four first columns indicate the proportion of cases, among 60 replicates, where each tested model was selected using AIC, given that data sets were generated under a particular model (*i*.*e*. true model). Column *d*AIC_true_ (*resp. d*AIC_wrong_) indicates the mean difference between the AIC of the model selected when the model selected is the true one (*resp*. when the model selected is not the true model) and the second best model (*resp*. being the true model or not).

| True Model | Selected Model | | | | $dAIC_{\text{true}}$ | $dAIC_{\text{wrong}}$ |
| --- | --- | --- | --- | --- | --- | --- |
| | $J_{\text{Exp}}$ | $J_{\text{Gauss}}$ | $J_{\text{ExpP}}$ | R.D. | | |
| $J_{\text{Exp}}$ | **0.50** | 0.32 | 0.05 | 0.13 | 0.90 | 1.10 |
| $J_{\text{Gauss}}$ | 0.25 | **0.45** | 0.02 | 0.28 | 1.30 | 0.67 |
| $J_{\text{ExpP}}$ | 0.17 | 0.05 | **0.78** | 0 | **217.50** | 0.76 |
| R.D. | 0.22 | 0.18 | 0 | **0.60** | 2.37 | 0.33 |

**Figure 2.**
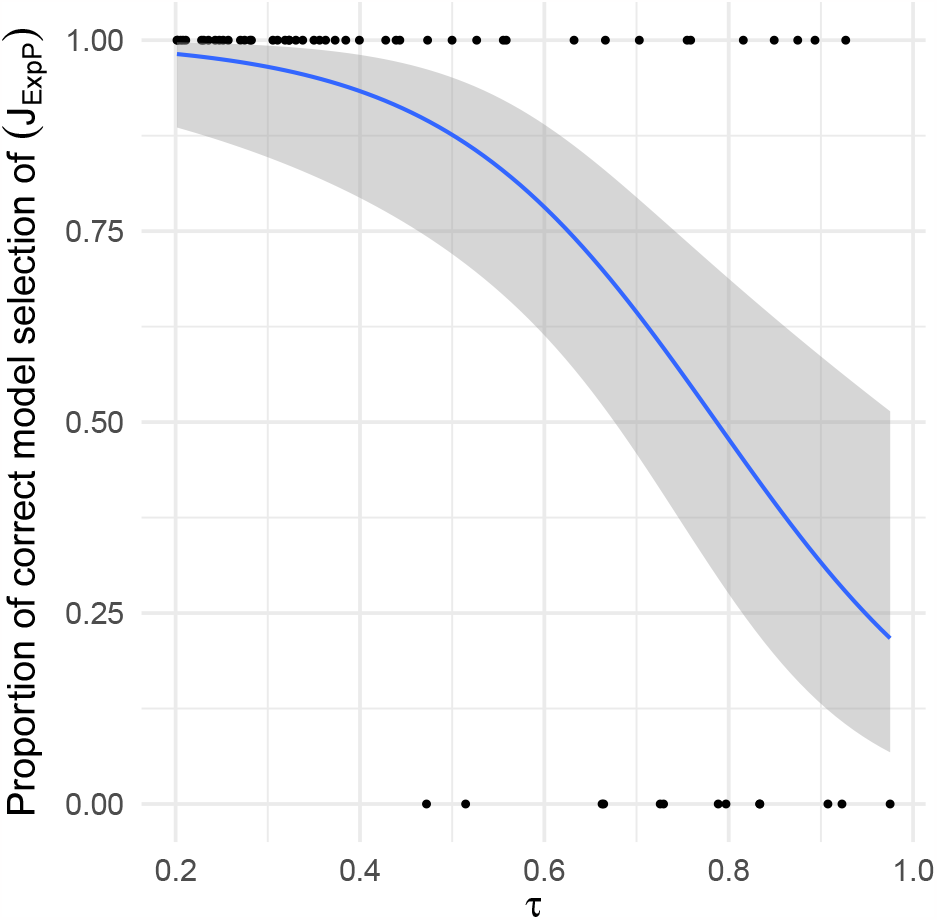
Logistic regression of the proportion of correct model selection of dispersal *J*_ExpP_ as a function of *τ*. Dots represent the values of *τ* used for the 60 replicates of simulated dispersal model *J*_ExpP_, and the estimated dispersal model (1 for a correct model selection of *J*_ExpP_ and 0 for a wrong model selection). The blue line corresponds to the predicted value of the proportion of correct model selection *J*_ExpP_ as a function of *τ*, and the grey area corresponds to the confidence envelope at 95%.

## 4. Case study: Invasion of poplar rust along the Durance River valley

### 4.1 Study site

We applied our approach to infer the dispersal of the plant pathogen fungus *Melampsora laricipopulina*, responsible for the poplar rust disease, from the monitoring of an epidemic invading the Durance River valley. Embanked in the French Alps, the Durance River valley constitutes a one-dimension ecological corridor that channels annual epidemics of the poplar rust pathogen *M. larici-populina* (Xhaard et al., 2012). Each year the fungus has to reproduce on larches (*Larix decidua*) that are located in the upstream part of the valley only. This constitutes the starting point of the annual epidemics. Then the fungus switches to poplar leaves and performs several rounds of infection until leaf-fall. Each infected leaf produces thousands of spores that are wind-dispersed. In our case study, *u*(*t, x*_*s*_) is the density of fungal infection at time *t* at point *x* on a poplar leaf. Each leaf has a carrying capacity of 750 fungal infections (Appendix S5).

All along the valley, the Durance River is bordered by a nearly continuous riparian forest of wild poplars (*Populus nigra*). The annual epidemic on poplars thus spreads downstream through the riparian stands, mimicking a one-dimension biological invasion (Xhaard et al., 2012). A previous genetic study showed that the epidemic was indeed initiated in an upstream location where poplars and larches coexist (Prelles), and progresses along the valley (Becheler et al., 2016). In autumn, the corridor is cleared for disease after leaf-fall. At 62 km downstream of the starting point of the epidemics, the Serre-Ponçon dam represents a shift point in the valley topology, with a steed-sided valley upstream and a larger riparian zone downstream. This delimitation led us to consider 2 values of growth rates *r* along the one-dimensional domain: *r*_up_ and *r*_dw_ (see Appendix S4 for details).

### 4.2 Monitoring of an annual epidemic wave

In 2008, rust incidence was monitored every three weeks from July to November at 12 sites evenly distributed along the valley (Figure 3). Sites were inspected during seven rounds of surveys. For a unique date (Oct. 22), the raw sampling was densified with 45 sites monitored instead of 12. We focused on young poplar trees (up to 2m high) growing on the stands by the riverside.

**Figure 3.**
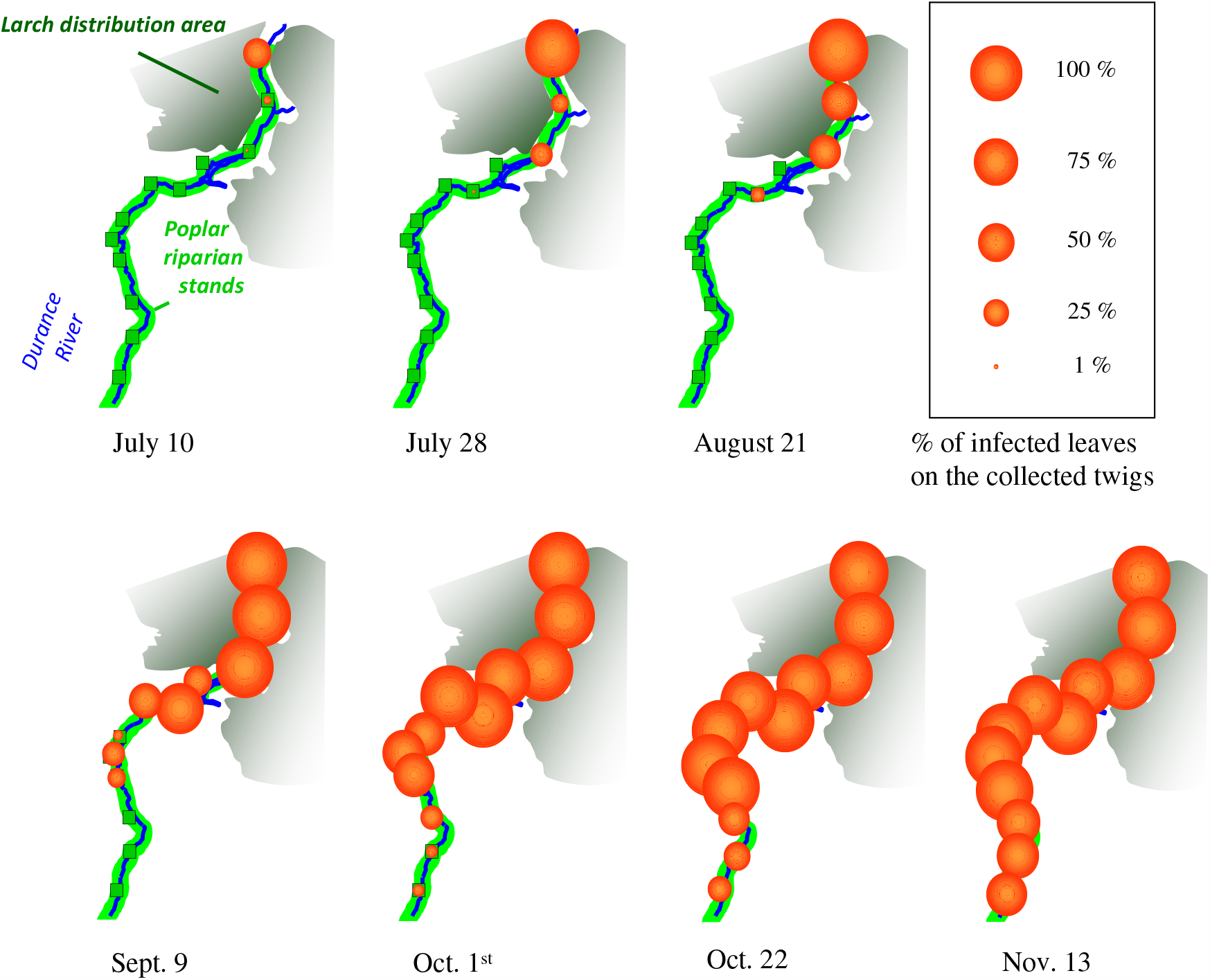
Poplar rust epidemic wave along the Durance River valley in 2008. The larch distribution area is represented in dark green, wild poplar riparian stands in pale green. The 12 study sites are represented by the green squares. Orange dots describe the evolution of the poplar rust epidemic through time (7 rounds of disease notation) and space (12 studied sites). Dot size is proportional to rust disease incidence assessed from the refined sampling.

Two monitorings were conducted, corresponding to the raw and refined sampling, as described in previous sections. For the raw sampling, we prospected each site at each date to search for rust disease by inspecting randomly distributed poplar trees (different trees at different dates for a given site). Depending on rust incidence and poplar tree accessibility, 40 to 150 trees (mean 74) were checked for disease. Each tree was inspected through a global scan of the leaves on different twigs until at least one infected leaf was found or after 30 s of inspection. The tree was denoted infected or healthy, respectively. This survey method amounts to minutely inspecting 10 leaves per tree, *i*.*e*. with the same efficiency of disease detection as through the refined sampling (see details of the statistical procedure in Appendix S3). The global scan procedure of the trees leads to equivalently surveying fewer and fewer leaves as the epidemic progresses. Optionally, when at least one tree was infected, and depending on available time, we carried out a refined sampling to collect more information on the variance in disease susceptibility (*i*.*e*. habitat suitability) among the sampling domain. The refined sampling consisted in randomly sampling 20 twigs on different trees and recording, for each, the total number of leaves and the number of infected leaves.

### 4.3 Dispersal and demographic processes ruling the epidemic wave

Model selection was used to decipher which dispersal process was best supported by the data set for five initial parameter values. The large AIC difference in favour of hypothesis *J*_ExpP_ indicates that poplar rust propagules assuredly disperse according to an exponential-power dispersal kernel along the Durance River valley (Table 3). Note that for all kernels, the five initial parameter values lead to similar estimations.

**Table 3:** Model selection for the epidemic of poplar rust along the Durance River valley. The Akaike information criteria are indicated for each model fitted to the real data set. The model best supported by the data is indicated in bold. AIC_median_ and AIC_sd_ represent the median and standard deviation among the AIC obtained from five initial parameter values.

| Dispersal | $AIC_{\text{median}}$ | $AIC_{\text{sd}}$ |
| --- | --- | --- |
| $J_{\text{Exp}}$ | 5461 | 5.81 |
| $J_{\text{Gauss}}$ | 5493 | 0.15 |
| $J_{\text{ExpP}}$ | **5163** | **0.01** |
| R.D. | 6190 | 0.03 |

The estimation of the parameters for the best model along with their confidence intervals (Appendix S4.3) are summarised in Table 4. The parameters of the Exponential-power kernel firstly indicate that the mean distance travelled by rust spores is estimated at 1.94 km. Second, its mean exponent parameter *τ* is 0.24. This value, much lower than 1, suggests substantial long-distance dispersal events. We also estimated the growth rates of the poplar rust epidemics along the Durance River valley. From upstream to downstream, their mean estimates are 0.085 and 0.023, respectively. The estimate of the parameter of the observation model, *γ*, is 4.82. This parameter represents how perceived probabilities of leaf infection differ among trees from true probabilities. The estimated value of 4.82 indicates some variability in the perception of infected leaves, but this variability is moderate because the shape of the underlying Beta-Binomial distribution approaches the Binomial distribution (for which perception differences are absent) (Figure 4, row 1). By contrast, the estimated value of the micro-scale fluctuation variance *σ*^2^ (1.29) suggests a substantial variability in leaf suitability between twigs. This is evidenced by comparing the shape of the estimated Gamma-Binomial distribution with a situation with negligible differences in receptivity between twigs (Figure 4, row 2, case *σ*^2^ = 0.01).

**Table 4:** Statistical summary of the inference of the parameters for the model best supported by the real data set *J*_ExpP_. We used the vector of parameters *θ* giving the lowest AIC value in the previous model selection procedure as initial parameter values of the R function mle2, to obtain maximum likelihood estimates of the vector of parameters 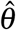 and of its matrix of variance-covariance 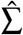. Summary statistics were derived from 1,000 random draws from the multivariate normal distribution with parameters 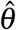 and 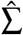 (see Appendix S4.3). Columns Estimate, *q−* 2.5% and *q−* 97.5% represent the estimated value of each parameter and the quantiles 2.5% and 97.5%, respectively.

| Parameter | Description | $q - 2.5\%$ | Estimate | $q - 97.5\%$ |
| --- | --- | --- | --- | --- |
| $r_{\text{up}}$ | Growth rate upstream | 0.0786 | 0.0853 | 0.0897 |
| $r_{\text{dw}}$ | Growth rate downstream | 0.0143 | 0.0230 | 0.0311 |
| $\lambda$ | Mean dispersal distance | 1.78 | 1.94 | 2.12 |
| $\tau$ | Kernel exponent | 0.226 | 0.243 | 0.260 |
| $\gamma$ | Tree perception | 3.49 | 4.82 | 6.11 |
| $\sigma^2$ | Variance in leaf suitability | 1.23 | 1.29 | 1.36 |

**Figure 4.**
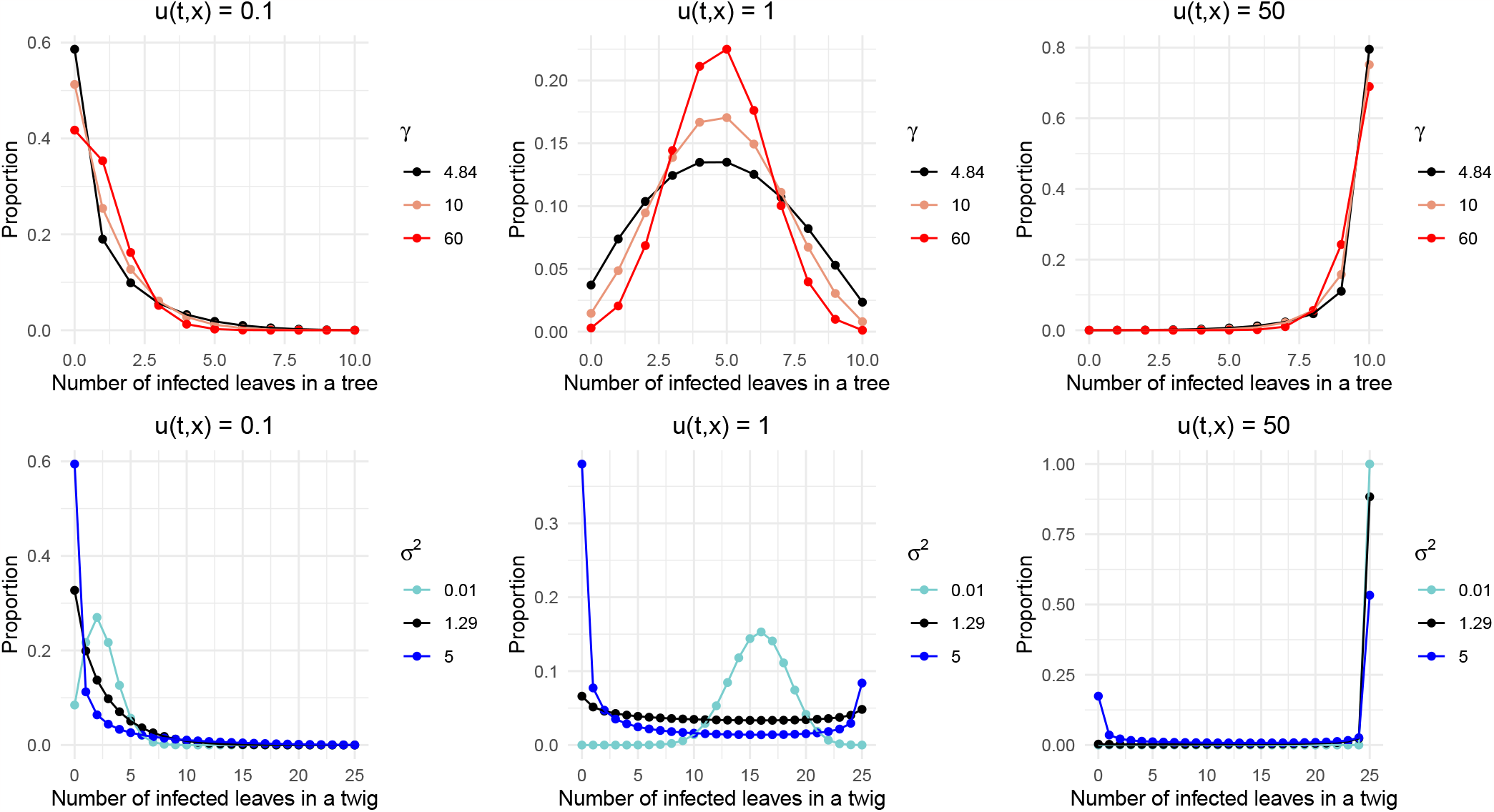
Distributions of the number of infected leaves in a tree and of the number of infected leaves in a twig, for increasing densities of infection *u*(*t, x*), and contrasted levels of environmental heterogeneity *σ* ^2^ and *γ*. The number of infected leaves in a tree follows a Beta-Binomial distribution (Eq. (S12)) with *σ* ^2^ = 1.29. Its density is plotted for three tree perceptions *γ*: 4.82 (estimated value on the real data set), 10 (intermediate value) and 60 for which the Beta-Binomial distribution is approaching a Binomial distribution. The number of infected leaves in a twig follows a Gamma-Binomial distribution (Eq. (S18)). Its density is plotted for three leaf suitabilities *σ* ^2^: 1.29 (estimated value on the real data set), 5 (a higher value) and 0.01 a value lowering variability in leaf suitability between twigs (when *σ* ^2^ tends to 0, all twigs share the same leaf suitability).

Model check consists in testing whether the selected model was indeed able –given the parameter values inferred above– to reproduce the observed data describing the epidemic wave that invaded the Durance River valley in 2008. To do so, we assessed the coverage rate of the raw sampling data (proportions of infected trees) based on their 95%-confidence intervals (Appendix S4.4, Figure 5). Over all sampling dates, the total coverage rate is high (0.71), which indicates that the model indeed captures a large part of the strong variability of the data. By comparison, coverage rates given by models *J*_Exp_ and *J*_Gauss_ (0.68 and 0.67, respectively) show a poorer fit to the data, especially for the first sampling date (Figures S6, S7) where the epidemic intensity is underestimated upstream and overestimated downstream.

**Figure 5.**
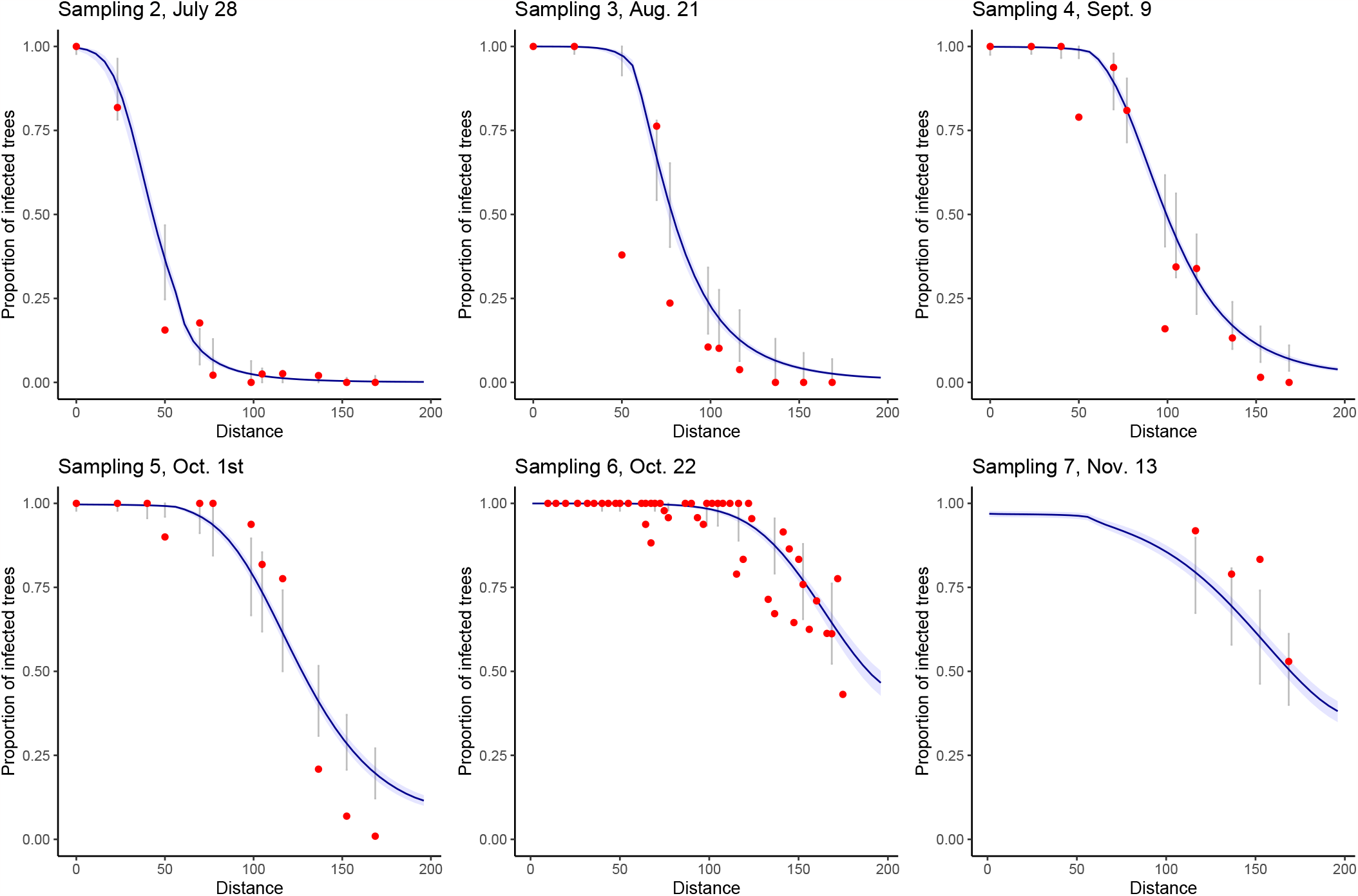
Model check under the selected dispersal model *J*_ExpP_: Coverage rates for the raw sampling. Each sampling date is represented on a separate graph. Sampling 1 is not represented because it corresponds to the initial condition of the epidemics for all simulations. Blue areas correspond to the pointwise 95% confidence envelopes for the proportion of infected trees, grey intervals correspond to the 95% prediction intervals at each site, *i*.*e*. taking into account the observation laws given the proportion of infected trees. Red points correspond to the observed data. Only four observations are available for sampling 7 because at this date (November 13) the leaves had already fallen from the trees located upstream the valley. The total coverage rate over all sampling dates is 0.71.

## 5 Discussion

This study combines mechanistic and statistical modelling to jointly infer the demographic and dispersal parameters underlying a biological invasion. A strength of the mechanistic model was to combine population growth with a large diversity of dispersal processes. The mechanistic model was coupled to a sound statistical model that considers different types of count data. These observation laws consider that habitat suitability and disease perception can vary over the sampling domain. Simulations were designed to prove that the demographic model can be confidently selected and its parameter values reliably inferred. Although the framework is generic, it was tuned to fit the annual spread of the poplar rust fungus *M. larici-populina* along the Durance River valley. This valley channels every year the spread of an epidemic along a one-dimensional corridor of nearly 200 km (Xhaard et al., 2012; Becheler et al., 2016). The monitoring we performed enables to build a comprehensive data set at a large spatial scale, which is mandatory to precisely infer the shape of the tail of dispersal kernels (Ferrandino, 1996; Kuparinen et al., 2007). A widely used alternative to the mechanistic-statistical approaches is to consider purely correlative approaches. However, the estimated parameters defining the strength of the temporal and spatial dependencies (as estimated for example using R-INLA package approach, Rue et al., 2009) will not allow to distinguish between the different shapes of dispersal kernels, which was the main goal of our work.

### 5.1 Estimation of the dispersal kernel of the poplar rust

This study provides the first reliable estimation of the dispersal kernel of the poplar rust fungus. Dispersal kernels are firstly defined by their scale, which can be taken to correspond to the mean dispersal distance. The mean dispersal distance obtained from the best model is 1.94 km with a 95% confidence interval ranging from 1.78 to 2.12 km. A non-systematic literature review identified only eight studies reporting dispersal kernels for plant pathogens that used data gathered in experimental designs extending over regions bigger than 1 km^2^ (Fabre et al., 2021). The mean dispersal distances of the four fungal pathosystems listed by these authors are 213 m for the ascospores of *Mycosphaerella fijiensis* (Rieux et al., 2014), 490 m for the ascospores of *Leptosphaeria maculans* (Bousset et al., 2015), 860 m for *Podosphaera plantaginis* (Soubeyrand et al., 2009a) and from 1380 to 2560 m for *Hymenoscyphus fraxineus* (Grosdidier et al., 2018). Our estimates for poplar rust are in the same range as the one obtained at regional scale for *Hymenoscyphus fraxineus*, the causal agent of Chalara ash dieback (Grosdidier et al., 2018).

Dispersal kernels can be further defined by their shape. We show that the spread of poplar rust is best described by a fat-tailed Exponential-power kernel. The “thin-tailed” kernels considered (Gaussian and exponential kernels) were clearly rejected by model selection. These results are in accordance with the high dispersal ability and the long-distance dispersal events evidenced in this species by population genetics analyses (Barrès et al., 2008; Becheler et al., 2016). Rust fungi are well-known to be wind dispersed over long distances (Brown and Hovmøller, 2002; Aylor, 2003). Recently, Severns et al. (2019) gathered experimental and simulation evidence that supports that wheat stripe rust spread supports theoretical scaling relationships from power law properties, another family of fat-tail dispersal kernel. In fact, many aerially dispersed pathogens are likely to display frequent long-distance flights as soon as their propagules (spores, insect vectors) escape from plant canopy into turbulent air layer (Ferrandino, 1993; Pan et al., 2010). Accordingly, four of these eight studies listed by Fabre et al. (2021) lent support to fat-tailed kernels, including plant pathogens as diverse as viruses, fungi, and oomycetes.

### 5.2 Effect of fat-tailed dispersal kernels on eco-evolutionary dynamics

The dynamics produced by the mechanistic integro-differencial models we use strongly depends on the tail of the dispersal kernel. Namely, when the equation is homogeneous (*i*.*e*. when the model parameters do not vary in space, leading to *r*(*x*) = *r*), it is well known that for any thin-tailed dispersal kernel *J* such that ∫_ℝ_ *J*(*z*)*e*^*λ*|*z*|^*dz <* +∞ for some *λ >* 0, the dynamics of *u*(*t, x*) is well explained using a particular solution called travelling wave. In this case, the invading front described by the solution *u*(*t, x*) moves at a constant speed (Aronson and Weinberger, 1978). By contrast, for a fat-tailed kernel, these particular solutions do not exist anymore, and the dynamic of *u*(*t, x*) describes an accelerated invasion process (Medlock and Kot, 2003; Garnier, 2011; Bouin et al., 2018). Here, we show that the dynamics of the poplar rust is better described as an accelerated invasion process rather than a front moving at a constant speed. Such accelerating wave at the epidemic front has been identified for several fungal plant pathogens dispersed by wind, including *Puccinia striiformis* and *Phytophthora infestans* the wheat stripe rust and the potato late blight, respectively (Mundt et al., 2009). However, it should be stated that fat-tailed kernels are not always associated with accelerated invasion processes. Indeed, fat-tailed kernels can be further distinguished depending on whether they are “regularly varying” (*e*.*g*. power law kernels) or “rapidly varying” (*e*.*g*. Exponential-power kernels) (Klein et al., 2006). Mathematically, it implies that power law kernels decrease even more slowly than any Exponential-power function. Biologically, fat-tailed Exponential-power kernels display rarer long-distance dispersal events than power law kernels. On the theoretical side, the kernel’s properties subtly interact with demographic mechanisms such as Allee effects to possibly cancel the acceleration of invasion. With weak Allee effects (*i*.*e*. the growth rate is density dependent but still positive), no acceleration occurs with rapidly varying kernels whereas an acceleration could be observed for some regularly varying kernels, depending on the strength of the density dependence (Alfaro and Coville, 2017; Bouin et al., 2021). For strong Allee effects (*i*.*e*. a negative growth rate at low density), no acceleration can be observed for all possible kernels (Chen, 1997). On the applied side, whether or not the epidemic wave is accelerating sharply impacts the control strategies of plant pathogens (Filipe et al., 2012; Ojiambo et al., 2015; Fabre et al., 2021).

### 5.3 Confidence in the inference of the dispersal process

The inference framework we developed is reasonably efficient in estimating the dispersal process with frequent long-distance dispersal events as generated by Exponential-power dispersal kernels. The numerical experiments clearly show that the lower the exponent parameter *τ* of the Exponential-power kernel, the higher the confidence in its selection.

Conversely, the identification of the dispersal process is less accurate with thin-tail kernels. The requirement for improving the capacity to distinguish between thin-tail kernels may lie in the sampling scheme. Here, our sampling sites are regularly spaced, over a large sampling domain of 200 km, which is better suited to monitor long-distance dispersal (Kuparinen et al., 2007). Sampling schemes with more frequent data in both time and space (or nested spatial sampling) might improve kernel identification.

We clearly observed that integro-differential models with Gaussian dispersal kernel on the one hand and reaction-diffusion equation on the other hand are well identified with our estimation procedure when the time and space sampling is dense enough. This result may at first appear striking as a common belief tends to consider that diffusion amounts to a Gaussian dispersal kernel. However, these two models represent different movement processes (Othmer et al., 1988). In addition, classical macroscopic diffusion, which is mainly based on Brownian motion (Othmer et al., 1988), often ignores the inherent variability among individuals’ capacity of movements and as a consequence does not accurately describe the dispersal at the population scale (Hapca et al., 2009). While it is reasonable to assume that a single individual disperses via Brownian motion, this assumption hardly extends to all individuals in the population. Accordingly, we believe that integro-differential models are better suited to take into account inter-individual behaviour as the dispersal kernel explicitly models the redistribution of individuals.

### 5.4 Robustness and portability of the method

A strength of the approach proposed is the detailed description of the observation laws in the statistical model. The derivation of their probability density functions allows to obtain an analytical expression of the likelihood function. Model inference was however not straightforward due to local optimum issues. In order to achieve satisfying computational efficiency, we developed an *ad hoc* hybrid strategy initiated from 20 initial values and combining the two classical Nelder-Mead and Nlminb optimisation algorithms. However, the framework of hierarchical statistical models (Cressie et al., 2009), whose inference is often facilitated by Bayesian approaches, could likely be mobilised to improve model fit. In particular, although the coverage rate of the tree sampling was correct, it could be further improved by relaxing some hypotheses. The orange-coloured uredinia being easily seen on green leaves, we assumed that the persons in charge of the sampling perfectly detect the disease as soon as a single uredinia is present on a leaf. However, even in this context, observation errors are likely present in our dataset as in any large spatio-temporal study. The latent variables used in hierarchical models are best suited to handle the fact that a tree observed to be healthy can actually be infected. False detection of infection could also be taken into account. This could make sense as a sister species, *M. alli-populina*, not easily discernible from *M. larici-populina* in the field, can also infect poplar leaves. This species can predominate locally in the downstream part of the Durance River valley. This could have led to over-estimate the disease severity at some locations. Yet, all infected leaves from twigs were collected and minutely inspected in the lab under a Stereo Microscope (25× magnification) to check for species identification.

More generally, the statistical part of the mechanistic-statistical approaches developed could be transposed to a wide range of organisms and sampling types. Sharing the sampling effort between raw and refined samples improves the estimations. The two distinct types of sampling (sampling of random leaves in trees, and of leaves grouped within twigs) apply to a wide range of species, which local distribution is aggregated into patches randomly scattered across a study site. Any biological study with two such distinct sampling types (as described in Figure 1) would fit the proposed statistical model. One can for example scale up the sampling by considering the plant (instead of the leaf) as the basic unit. Moreover, the framework naturally copes with the diversity of sampling schemes on the ground such as the absence of one sample type for all or part of the sampled sites and dates. Finally, we used the first sampling date to estimate independently the initial population densities *u*(0, *x*) that were then fixed among all simulated epidemics. Future works could as well jointly estimate *u*(0, *x*) as part of *θ* .

The mechanistic part of the model could also handle a wider diversity of hypotheses. First, the model can be adapted to take into account a wider range of dispersal kernels, such as regularly varying kernels (see above). Second, the model can also easily be adapted to take into account parameter heterogeneity in time and space of its parameters. Similarly, one may easily assume that the growth rate depends on daily meteorological variables. Finally, we ignore the influence of the local fluctuations of the population size on the macro-scale density of the population when stochastic fluctuations can influence epidemic dynamics (Rohani et al., 2002). Here, we neglect this influence by considering that the average population size is relevant when habitat units are aggregated. Relaxing this hypothesis could be achieved by incorporating stochastic integro-differential equations. The inference of such models is currently a front of research.

### 5.5 Future directions

As biological invasions are regularly observed retrospectively, carrying out spatio-temporal monitoring is often highly difficult, when possible. A small number of longitudinal temporal data makes model inference very difficult, in particular for its propensity to properly disentangle the effect of growth rate and dispersal. Incorporating genetic data into the framework proposed here is a challenge that must be met to get around this problem. Indeed, colonisation and demographic effects such as Allee effect generate their own specific genetic signatures (Dennis, 1989; Lewis and Kareiva, 1993; Miller et al., 2020). Similarly, genetic data could help to identify the dispersal kernel underlying the invasion process, as the population will exhibit an erosion of its neutral diversity with a thin-tailed kernel (Edmonds et al., 2004; Hallatschek et al., 2007). Conversely, genetic diversity can be preserved all along the invasion front with a fat-tailed kernel, because of the long-distance dispersal of individuals from the back of the front, where genetic diversity is conserved (Fayard et al., 2009; Bonnefon et al., 2014).

## Acknowledgements

We warmly thank all the collectors who participated in the monitoring of the poplar rust disease spread along the Durance River valley: Audrey Andanson, Béranger Bertin, Olivier Caël, Bénédicte Fabre, Christine Gehin, Claude Husson, Benôıt Marçais, and Agathe Vialle. We also thank Benôıt Marçais for fruitful discussions on disease monitoring, Bénédicte Fabre for the calculus of the density in uredinia on a poplar leaf, and Fabrice Elegbede for advices on statistical analyses. This work was supported by grants from the French National Research Agency (ANR-09-BLAN-0145, EMILE project; ANR-18-CE32-0001, CLONIX2D project; ANR-14-CE25-0013, project NONLOCAL, ANR-11-LABX-0002-01, Cluster of Excellence ARBRE; 20-PCPA-0002, BEYOND project). Constance Xhaard was supported by a PhD fellowship from the French Ministry of Education and Research (MESR) and by Postdoc fellowship from the French National Research Agency (ANR-09-BLAN-0145, EMILE project) . Méline Saubin was supported by a PhD fellowship from INRAE and the French National Research Agency (ANR-18-CE32-0001, CLONIX2D project).

## Author contributions

Constance Xhaard, Pascal Frey, and Fabien Halkett supervised the disease monitoring. Jérôme Coville, Frédéric Fabre, Fabien Halkett, and Samuel Soubeyrand conceived and designed the study. Jérôme Coville provided a mathematical expertise on modelling long-range dispersal as well as codes of simulation for the mechanistic models. Samuel Soubeyrand established the observation laws. Frédéric Fabre supervised the statistical analyses. Constance Xhaard and Fabien Halkett did preliminary analyses. Méline Saubin updated the code and did the statistical analyses. Jérôme Coville, Frédéric Fabre, Fabien Halkett, Méline Saubin, and Samuel Soubeyrand contributed to the writing of the manuscript. All authors read and approved the manuscript.

## Data accessibility

R and C++ scripts for model simulations and statistical analyses, as well as count data for the biological application, are available on a public Zenodo repository (DOI:10.5281/zenodo.7906840), extracted from a public GitLab repository: https://gitlab.com/saubin.meline/mechanistic-statistical-model. 10.5281/zenodo.7906840

