## Supplementary Material for "A mechanistic-statistical approach to infer dispersal and demography from invasion dynamics, applied to a plant pathogen"

[#] These authors co-directed this work.

**Corresponding authors:**

### S1 Numerical scheme

We use an implicit Euler scheme combined with a finite difference scheme (see Allaire, 2005 for details) to compute the solution $u(t,x)$ of the reaction-diffusion equation over $[-R,R]\times[0,T]$, with $2\times R$ the length of the modelled environment, and $T$ the duration of the modelled process. For the integro-differential equation, we use an explicit Euler scheme. More precisely, we perform a standard explicit Euler time discretisation of the equation:

$$\frac{\partial u}{\partial t}(t,x) \approx \frac{u(t+\delta,x)-u(t,x)}{\delta} \tag{S1}$$

that leads to:

$$\begin{aligned} u(t_{n+1},x) = u(t_n,x) + \delta\left(\int_{-R}^{R} J(x-y)[u(t_n,y)-u(t_n,x)]\,dy\right) \\ + \delta r(x) u(t_n,x)\left(1-\frac{u(t_n,x)}{K}\right) \end{aligned} \tag{S2}$$

where $\{t_n = n\delta = nT/N : n = 0,\ldots,N\}$ is a series of increasing times separated by $\delta = T/N > 0$, and $N$ is the number of time steps in the series. For the space discretisation, we define a regular grid $\{x_i = -R+i\varepsilon = -R+2Ri/I : i = 0,\ldots,I\}$ with $I+1$ points separated by $\varepsilon = 2R/I > 0$. We make the following approximation for all $x$ in $[-R,R]$:

$$u(t_n,x) \approx \sum_{i=0}^{I} u(t_n,x_i)\mathbb{1}_{[x_i,x_i+\varepsilon)}(x) \tag{S3}$$

where $x \mapsto \mathbb{1}_{[x_i,x_i+\varepsilon)}(x)$ is the indicator function that gives 1 if $x\in[x_i,x_i+\varepsilon)$, 0 otherwise. Based on this approximation, we only need to compute $u(t,x)$ at points $x_i$, $i = 0,\ldots,I$. Plugging Approximation (S3) in the integral of Equation (S2) computed for $x = x_i$ yields:

$$\begin{aligned}
&\int_{-R}^{R} J(x_i - y)[u(t_n,y) - u(t_n,x_i)]\,dy \\
&\qquad \approx \int_{-R}^{R} J(x_i - y)\left[\left(\sum_{j=0}^{I} u(t_n,x_j)\mathbb{1}_{[x_j,x_j+\varepsilon)}(y)\right) - u(t_n,x_i)\right] dy \\
&\qquad = \left(\sum_{j=0}^{I} u(t_n,x_j)\int_{-R}^{R} J(x_i - y)\mathbb{1}_{[x_j,x_j+\varepsilon]}(y)\,dy\right) - \left(u(t_n,x_i)\int_{-R}^{R} J(x_i - y)dy\right) \\
&\qquad \approx \varepsilon\left(\sum_{j=0}^{I} u(t_n,x_j)J(x_i - x_j)\right) - \varepsilon\, u(t_n,x_i)\sum_{j=0}^{I} J(x_i - x_j)
\end{aligned} \tag{S4}$$

Let us define the matrix $\mathbf{J}^{in} := (J(x_i - x_j))_{0\le i,j\le I}$ whose element $(i,j)$ is $\mathbf{J}^{in}_{ij} = J(x_i - x_j)$. We get the following numerical scheme:

$$\begin{aligned}
u(t_{n+1},x_i) = u(t_n,x_i) + \delta\varepsilon\left[\sum_{j=0}^{I}\mathbf{J}^{in}_{ij}u(t_n,x_j) - u(t_n,x_i)\left(\sum_{j=0}^{I}\mathbf{J}^{in}_{ij}\right)\right] \\
+ \delta r(x_i)u(t_n,x_i)\left[1 - \frac{u(t_n,x_i)}{K}\right]
\end{aligned} \tag{S5}$$

By defining the vectors $\mathbf{U}(t_n) = (u(t_n,x_i))_{0\le i\le I}$, $\mathbf{R} = (r(x_i))_{0\le i\le I}$ and $\mathbf{1} = (1)_{0\le i\le I}$, we have to solve the linear system:

$$\mathbf{U}(t_{n+1}) = \mathbf{U}(t_n) + \delta\varepsilon\left\{\mathbf{J}^{in}\mathbf{U}(t_n) - \mathbf{U}(t_n)\cdot(\mathbf{J}^{in}\mathbf{1})\right\} + \delta\{\mathbf{R}\cdot\mathbf{U}(t_n)\}\cdot\left\{\left(1 - \frac{\mathbf{U}(t_n)}{K}\right)\right\} \tag{S6}$$

where $\cdot$ is the element-wise multiplication operator.

### S2 Distributions of the population measurements

#### S2.1 Term designations for the sampling units

In our biological application, a poplar leaf represents a habitat unit, a twig represents a group of habitat units, and a tree represents a habitat bloc. For clarity, we refer to leaves, twigs and trees in the following explanations. We call a sampling site a surveyed area along the valley, containing several hundreds of trees. Further adaptations of this model to other sampling units would only require adapting this initial vocabulary (Figure 1).

#### S2.2 Raw sampling

In the raw sampling, trees represent the sampling units, and $B_{st}$ trees are observed in site $s$ at time $t$. For each tree $b \in \{1,\ldots,B_{st}\}$, we measure the presence/absence of the pathogen by monitoring an equivalent number of $M$ leaves within $b$ (see Appendix S3 below for the determination of $M$). A tree is infected if at least one pathogen lesion has been detected, in at least one leaf of the tree. The observation in site $s$ at time $t$ is the number $Y_{st}$ of infected trees.

Now, let us derive the probabilistic law of the presence/absence of the pathogen in any tree $b$ observed in site $s$ at time $t$. In this paragraph, subscripts $s$, $t$, and $b$ are generally omitted to avoid cumbersome notation. We first remind that the numbers of pathogen lesions $N_i(t)$ in the leaf $i \in \{1,\ldots,M\}$ observed in tree $b$, given $R_i(t)$ and $u(t,x_s)$, are independent and Poisson distributed (see Eq. (2) in the main text):

$$N_i(t) \mid u(t,x_s), R_i(t) \underset{\text{indep.}}{\sim} \text{Poisson}(u(t,x_s)R_i(t)) \tag{S7}$$

In the raw sampling, $M$ leaves are sampled at different locations on the tree (*i.e.* they belong to different groups, referred to as twigs), but further information about the twigs is not known. Thus, in the following, we take into account the twig structure without exploiting twig information. The leaves of a given twig $g$ on tree $b$ share at time $t$ the same suitability $\mathcal{R}_g(t)$, which is unobserved and Gamma distributed like in Eq. (3) in the main text (for all leaves $i$ in twig $g$, $R_i(t) = \mathcal{R}_g(t)$). Given the suitabilities $\{\mathcal{R}_g(t) : g = 1,\dots,G\}$ of twigs which compose tree $b$ and given the absence of data about the twigs, $R_i(t)$ ($i \in \{1,\dots,M\}$) are independent and identically distributed under the discrete empirical probability distribution:

$$\hat{F}_G(r) = \frac{1}{G}\sum_{g=1}^{G} \mathbb{1}(r \leq \mathcal{R}_g(t)) \tag{S8}$$

where $\mathbb{1}(\cdot)$ is the indicator function. Therefore, $N_i(t)$ ($i \in \{1,\dots,M\}$) given $\{\mathcal{R}_g(t) : g = 1,\dots,G\}$ and $u(t,x_s)$ are independent and their probability distribution is, using Eqs. (S7)–(S8):

$$P[N_i(t) = n \mid u(t,x_s), \{\mathcal{R}_g(t) : g = 1,\dots,G\}] = \frac{1}{G}\sum_{g=1}^{G} \exp(-u(t,x_s)\mathcal{R}_g(t))\frac{(u(t,x_s)\mathcal{R}_g(t))^n}{n!} \tag{S9}$$

The suitability $\mathcal{R}_g(t)$ being Gamma distributed with shape and scale parameters $\sigma^{-2}$ and $\sigma^2$, respectively, the right-hand-side of Eq. (S9) is a Monte Carlo approximation of the integral:

$$\begin{aligned}\int_{\mathbb{R}_+} \exp(-u(t,x_s)r)\frac{(u(t,x_s)r)^n}{n!}\frac{1}{(\sigma^2)^{\sigma^{-2}}\Gamma(\sigma^{-2})}r^{\sigma^{-2}-1}e^{-r/\sigma^2}dr \\ = \frac{\Gamma(n+\sigma^{-2})}{(n!)\Gamma(\sigma^{-2})}\left(1-\frac{u(t,x_s)}{u(t,x_s)+\sigma^{-2}}\right)^{\sigma^{-2}}\left(\frac{u(t,x_s)}{u(t,x_s)+\sigma^{-2}}\right)^n\end{aligned} \tag{S10}$$

which coincides with the probability distribution of the Negative–Binomial law (*i.e.* the Gamma-Poisson mixture distribution) given by Eq. (4) in the main text. The larger $G$, the more precise the approximation. Consequently, $N_i(t)$ ($i \in \{1,\ldots,M\}$) given $u(t,x_s)$ are asymptotically independent and distributed under the Negative–Binomial distribution given by Eq. (4) in the main text. Based on this approximation, the infections of leaves from tree $b$ in site $s$ at time $t$ are asymptotically independent and distributed under Bernoulli distributions with success probability:

$$
\begin{aligned}
p_{st}^{\text{leaf}} &= P(N_i(t) > 0 \mid u(t,x_s)) \\
&= 1 - P(N_i(t) = 0 \mid u(t,x_s)) \\
&= 1 - (1 + u(t,x_s)\sigma^2)^{-1/\sigma^2}
\end{aligned} \tag{S11}
$$

The people who carried out the sampling observed a number $M$ of leaves on tree $b$. Due to the particular configuration of the foliage of each tree, we assumed that the number $Y_{stb}^{\text{leaf}}$ of infected leaves among the $M$ leaves observed in tree $b$ is approximately distributed under a Beta-Binomial distribution with mean $Mp_{st}^{\text{leaf}}$ and tree perception parameter $\gamma$:

$$
Y_{stb}^{\text{leaf}} \mid u(t,x_s) \sim_{approx.} \text{Beta-Binomial}(M, p_{st}^{\text{leaf}}, \gamma) \tag{S12}
$$

Accordingly, the probability, as *perceived* by people in charge of the sampling, of leaf infection on the set of $M$ leaves observed on a given tree, is distributed according to a Beta distribution. The Beta distribution is centred around the true probability of leaf infection $p_{st}^{\text{leaf}}$ and allows *perceived* probability to vary from tree to tree depending on the tree perception parameter $\gamma$. It follows that the infection of tree $b$ is approximately distributed under the Bernoulli distribution with success probability:

$$\begin{aligned} p_{st}^{\text{tree}} &= P(Y_{stb}^{\text{leaf}} > 0 \mid u(t, x_s)) \\ &= 1 - P(Y_{stb}^{\text{leaf}} = 0 \mid u(t, x_s)) \\ &= 1 - \frac{\text{Beta}[\gamma p_{st}^{\text{leaf}}, M + \gamma(1 - p_{st}^{\text{leaf}})]}{\text{Beta}[\gamma p_{st}^{\text{leaf}}, \gamma(1 - p_{st}^{\text{leaf}})]} \end{aligned} \tag{S13}$$

where $p_{st}^{\text{leaf}}$ is given by S11 and Beta represents the beta function. It follows that the probability distribution functions of the number $Y_{st}^{\text{tree}}$ of infected trees infected among the $B_{st}$ trees observed satisfy, for all sampling sites $s$ and sampling times $t$:

$$\begin{aligned} f_{st}^{\text{raw}}(y) &= P[Y_{st}^{\text{tree}} = y \mid u(t, x_s)] \\ &= f_{\text{Binomial}(B_{st}, p_{st}^{\text{tree}})}(y) \end{aligned} \tag{S14}$$

where $f_{Binomial}$ is the density of the Binomial distribution.

#### S2.3 Refined sampling

In the refined sampling, $G_{st}$ twigs (*i.e.* groups of spatially connected leaves) are sampled in site $s$ at time $t$. Here, the twig information (the number of twigs and the distribution of leaves on twigs) are known but the suitability $\mathcal{R}_g(t)$ of leaves in a twig $g$ remains unobserved. The numbers of pathogen lesions $N_i(t)$ in the observed leaves $i \in \{1, \ldots, M_{stg}\}$ of twig $g$ given $\mathcal{R}_g(t)$ and $u(t, x_s)$ are independent and Poisson distributed:

$$N_i(t) \mid u(t, x_s), \mathcal{R}_g(t) \underset{\text{indep.}}{\sim} \text{Poisson}(u(t, x_s)\mathcal{R}_g(t)) \tag{S15}$$

Then, the numbers of infected leaves $Y_{stg}^{\text{leaf}}$ (*i.e.* leaves with at least one pathogen lesion) given $\mathcal{R}_g(t)$ and $u(t,x_s)$ are independent and distributed under the following Binomial distributions:

$$Y_{stg}^{\text{leaf}} \mid u(t,x_s), \mathcal{R}_g(t) \underset{\text{indep.}}{\sim} \text{Binomial}(M_{stg}, 1 - e^{-u(t,x_s)\mathcal{R}_g(t)}) \tag{S16}$$

In addition,

$$u(t,x_s)\mathcal{R}_g(t) \mid u(t,x_s) \underset{\text{indep.}}{\sim} \text{Gamma}(\sigma^{-2}, u(t,x_s)\sigma^2) \tag{S17}$$

Using Eqs. (S15)–(S17), $Y_{stg}^{\text{leaf}}$ given $u(t,x_s)$ are independent and follow Gamma-Binomial mixture distributions:

$$\begin{aligned} f_{st}^{\text{ref}}(y) &= P[Y_{stg}^{\text{leaf}} = y \mid u(t,x_s)] \\ &= \int_0^\infty f_{\text{Binomial}(M_{stg},1-e^{-z})}(y) f_{\text{Gamma}(\sigma^{-2},u(t,x_s)\sigma^2)}(z) dz \end{aligned} \tag{S18}$$

where $f_{\text{Gamma}}$ is the density of the Gamma distribution. Note that this Gamma-Binomial mixture distribution is an over-dispersed Binomial distribution like the Beta-Binomial distribution.

### S3 Estimation of the number of leaves efficiently observed during tree scans

A problem inherent to the raw sampling design is that we do not know the number of leaves observed during the scan of the trees, contrary to the twig data for which we counted both the number of infected leaves and the total number of leaves carried by each observed twig. In other words, an inspected tree is a set of leaves of unknown size.

We assumes in Eq. (S12) that the number $Y_{stb}^{\mathrm{leaf}}$ of infected leaves among the $M$ leaves observed in tree $b$ is approximately distributed under a Beta-Binomial distribution with mean $Mp_{st}^{\mathrm{leaf}}$ and tree perception parameter $\gamma$. Parameter $\gamma$ is however an unknown parameter. To overcome this parameter when calculating the average number of leaves observed per tree, we use the fact that on average the number of infected leaves is the same with a binomial distribution:

$$Y_{stb}^{\mathrm{leaf}} \mid u(t, x_s) \sim_{approx.} \mathrm{Binomial}(M, p_{st}^{\mathrm{leaf}}) \tag{S19}$$

From this distribution, we obtain at each site $s$ and date $t$ the probability $p_{st}^{\mathrm{tree}}$ that a tree is infected as a function of both the probability $p_{st}^{\mathrm{leaf}}$ that a leaf is infected and the number $M$ of leaves observed on a tree:

$$
\begin{aligned}
p_{st}^{\text{tree}} &= P(Y_{stb}^{\text{leaf}} > 0 \mid u(t, x_s)) \\
&= 1 - P(Y_{stb}^{\text{leaf}} = 0 \mid u(t, x_s)) \\
&= 1 - (1 - p_{st}^{\text{leaf}})^M
\end{aligned}
\tag{S20}
$$

Thus, the number of leaves on a tree satisfies:

$$
M = \frac{\log(1 - p_{st}^{\text{tree}})}{\log(1 - p_{st}^{\text{leaf}})} \tag{S21}
$$

Let us use as approximations of $p_{st}^{\text{tree}}$ the observed proportions $q_{st}^{\text{tree}}$ of infected trees at sites $s$ and dates $t$, and as approximations of $p_{st}^{\text{leaf}}$ the observed proportions $q_{st}^{\text{leaf}}$ of infected leaves (calculated from twig data). Then, an estimate $\hat{\lambda}_M$ of the mean number of leaves $\lambda_M$ by tree is given by:

$$
\hat{\lambda}_M = \text{round}\left(\frac{1}{N}\sum_{i=1}^{N} \frac{\log(1 - q_{st}^{\text{tree}})}{\log(1 - q_{st}^{\text{leaf}})}\right) \tag{S22}
$$

with $N$ the number of pairs $(s,t)$ (*i.e.* sampling sites and dates) displaying both tree and twig data. Proportions of infection $q_{st}^{\text{tree}} = 1$ and $q_{st}^{\text{leaf}} = 1$ where approximated to $1 - 10^{-16}$ for numerical considerations. This procedure led to $\hat{\lambda}_M = 10$. This value may appear low. However, $\lambda_M$ does not correspond to the actual mean number of leaves carried by an entire young tree but amounts to the mean number of leaves effectively inspected during tree scan, *i.e.* those observed as minutely as for the twig data in a limited time (see Eq. (S13)). It is important to note that for each tree the tree scan stops when an infected leaf is observed, or after 30 s of inspection. Therefore, the number of inspected leaves per tree can be very low in highly infected sites.

For the practical identifiability studies, we set $\lambda_M = 10$. For parameter inference on the real data set a different value of $(\hat{\lambda}_M)_t$ was estimated for each sampling date, from the observed proportions $q_{st}^{\text{tree}}$ of infected trees and the observed proportions $q_{st}^{\text{leaf}}$ of infected leaves at date $t$ (Table S1).

Table S1: Estimated number of leaves effectively observed per tree for each sampling date $t$, $(\hat{\lambda}_M)_t$. The values of $(\hat{\lambda}_M)_t$ were used in the application on the real data set.

| Date t | $(\hat{\lambda}_M)_t$ |
|---|---|
| 1 | 40 |
| 2 | 24 |
| 3 | 6 |
| 4 | 3 |
| 5 | 5 |
| 6 | 1 |

### S4 Simulation details

Computations were performed with the `R` software environment (R Core Team, 2018). The vector of initial population densities $u(0,x)$ for $x$ over $[-R,R]$ was estimated from the data of the first sampling date, by fitting a general model for analysis of dose-response data (package `Drc` on `R`, Ritz et al., 2015). This vector of initial population densities represented the initial condition of all simulations. We modelled $N = 1500$ time steps and $I = 400$ points in space. Because of the numerical scheme, with these parameters the reaction-diffusion dispersal model R.D. required an upper limit for parameter $\lambda$: we set $\lambda_{up} = 23$ for this model.

To fit our real case study, for all simulations we set $R = 100$ km, for a 200 km long river valley, and the epidemic was monitored over $T = 150$ days. We considered a shift in the environment topology at $d = 0.31\%$ of the valley, which corresponds to the delimitation observed in the Durance River valley with the Serre-Ponçon dam at 62 km downstream of the starting point of the epidemic. Therefore, for all simulations, the two growth rates $r_{up}$ and $r_{dw}$ apply to continuous segments of proportions $d$ and $1-d$ of the monitored space, respectively.

#### S4.1 Practical parameter identifiability

Simulations were performed as follows in three steps.

**Step 1** : Simulation of a realistic epidemic. Given a hypothetical dispersal model ($J_{\text{Exp}}$, $J_{\text{Gauss}}$, $J_{\text{ExpP}}$ or R.D.), values in the parameter vector $\theta = (\theta_r, \theta_J, \gamma, \sigma^2)$ are independently and randomly drawn from dedicated distributions encompassing a large diversity of invading scenarios and specified in Table S1. We then simulate the corresponding epidemic along the 1D spatial domain $[-R,R]$. This epidemic is considered 'realistic' if a set of requirements on the observed proportion of infected trees $P_{s,t}$ on the farther downstream site ($s = R$) is met:

- $P_{R,30} < 0.1$ (the proportion of infected trees after one month is lower than 10%);
- $P_{R,75} < 0.5$ (the proportion of infected trees after two and a half months is lower than 50%);
- $P_{R,150} > 0.1$ (the proportion of infected trees after five months is higher than 10%);
- $P_{R,150} < 0.8$ (the proportion of infected trees after five months is lower than 80%).

Step 1 is complete once a candidate vector $\theta$ leads to an epidemic satisfying the four conditions described above (*i.e.* the simulation of $\theta$ and the epidemic is repeated while the four conditions are not satisfied). Thereafter, the vector finally retained in Step 1 is denoted $\theta_{\text{true}}$.

Table S1: Marginal distributions used to randomly sample the model parameters included in $\theta = (\theta_r, \theta_J, \gamma, \sigma^2)$ before checking the requirements detailed in Step 1, with $\theta_r = (r_{\text{dw}}, \omega)$ and $\theta_J = (\lambda)$ or $\theta_J = (\lambda, \tau)$ depending on the model.

| Parameter | Distribution | Interval |
|---|---|---|
| $r_{\text{dw}}$ | Log-Uniform | [0.01, 0.5] |
| $\omega$ | Uniform | [-2, 4] |
| $\lambda$ | Log-Uniform | [0.2,5] |
| $\tau$ | Log-Uniform | [0.2,1] |
| $\gamma$ | Log-Uniform | [2,20] |
| $\sigma^2$ | Log-Uniform | [0.01, 15] |

**Step 2** : Simulation of the sampling process. We consider a sampling design similar to our real experiment with six sampling dates and 12 sampling locations regularly spread over 150 days and 200 km, respectively ($R = 100$ km). As for our real data, we increase the location density for the fifth date, with 45 locations instead of 12. For each date and location, the raw sampling consists in simulating the observed sanitary status of 10 leaves per tree from 100 trees, and the refined sampling consists in simulating the observed sanitary status of 25 spatially connected leaves from 20 twigs,

the simulations being performed given $\theta_{\text{true}}$. The resulting data set is denoted $\mathcal{D}_{\text{true}}$.

**Step 3** : Parameter estimation. We use the data $\mathcal{D}_{\text{true}}$ to estimate the model parameters by minimizing the logarithm of the likelihood function $L(\theta)$. In our case, preliminary tests revealed that classical optimisation algorithms were not accurate enough to provide satisfactory rates of convergence due to local optimum problems. Thus, we adopt a hybrid strategy combining first a Nelder-Mead algorithm (improving global search ability) and then a Nlminb algorithm (for its high computational efficiency). Specifically, we proceed in three substeps described below, the crucial stage consisting in finding initial values that give a satisfactory rate of convergence.

**Step 3.1** : Using Step 1, we generate 500 vectors $\theta_{\text{init}}$. Note that this step was only performed once for all the estimations performed in this article. We provide in Figure S1 a comparison of the initial distribution of parameters as stated in Table S1, and of the distribution of parameters in the vector $\theta_{\text{init}}$, *i.e.* leading to "realistic" epidemics.

**Step 3.2** : The corresponding 500 likelihood values $L(\theta_{\text{init}})$ are calculated given $\mathcal{D}_{\text{true}}$. Then, the 20 vectors $\theta_{\text{init}}$ corresponding to the 20 largest likelihood values are used as initial values for 50 steps of a NELDER-MEAD optimisation routine (`R` function `optim`), resulting in 20 updated initial parameter vectors $\theta_{\text{init2}}$ depending on $\mathcal{D}_{\text{true}}$. The new initial vectors $\theta_{\text{init2}}$ that do not satisfy lower bounds $\theta_{\text{low}}$ and upper bounds $\theta_{\text{up}}$ are excluded. We used $\theta_{\text{low}} = (r_{\text{dw}} = 0.001, \omega = -7, \lambda = 0.02, \tau = 0.02, \gamma = 1.05, \sigma^2 = 10^{-7})$ and $\theta_{\text{up}} = (r_{\text{dw}} = 0.5, \omega = 4, \lambda = 10, \tau = 1, \gamma = 30, \sigma^2 = 20)$, with $\lambda = 23$ in $\theta_{\text{up}}$ instead of 10 for the R.D. model. The validity intervals defined by $\theta_{\text{low}}$ and $\theta_{\text{up}}$ encompass the intervals used to simulate $\theta$ (see Table S1). The likelihood values of the $n_{\text{init}}$

remaining vectors $L(\theta_{\text{init2}})$ are calculated (given $\mathcal{D}_{\text{true}}$) and ranked in descending order.

**Step 3.3** : $\theta$ is then estimated using the NLMINB optimisation routine with lower and upper bounds $\theta_{\text{low}}$ and $\theta_{\text{up}}$, respectively. The initial parameter values are set to the first vector $\theta_{\text{init2}}$ as ordered in the previous step. The estimated parameter values, say $\theta_{\text{estim}}$, are accepted if the `nlminb` function in `R` delivered a successful convergence diagnostic (with tunning parameters `rel.tol`=$5.10^{-5}$ and `iter.max`=3000). If not, the second vector $\theta_{\text{init2}}$ is used, and so on until reaching convergence or testing the $n_{\text{init}}$ initial vector's values selected at step 3.2. In the latter case, a convergence failure is obtained. Overall, this algorithm allows to obtain high rates of convergence.

These three steps were reiterated until deriving the estimation of $n = 160$ realistic epidemics for each dispersal model. Checking for practical identifiability of parameters basically relies on plotting for each dispersal model the cloud of points between $\theta_{\text{true}}$ and $\theta_{\text{estim}}$ (Figures S2, S3, S4, S5) and computing the corresponding correlations. Among all simulations performed, the proportions of convergence were 0.98 for dispersal $J_{\text{Exp}}$, $J_{\text{Gauss}}$, and R.D. and 0.99 for $J_{\text{ExpP}}$. A simulation converged when the convergence diagnostic of the algorithm indicated a convergence, and when all parameters were estimated inside intervals defined by $\theta_{\text{low}}$ and $\theta_{\text{up}}$. In the small number of simulations where the value of $\lambda_{\text{estim}}$ proposed by the optimisation algorithm was higher than 23 (which is the upper limit of our numerical scheme, Appendix S1), the simulation was still considered convergent with $\lambda_{\text{estim}} = 23$. This configuration can occur in particular when trying to fit dispersal R.D. on datasets simulated according to $J_{\text{ExpP}}$.

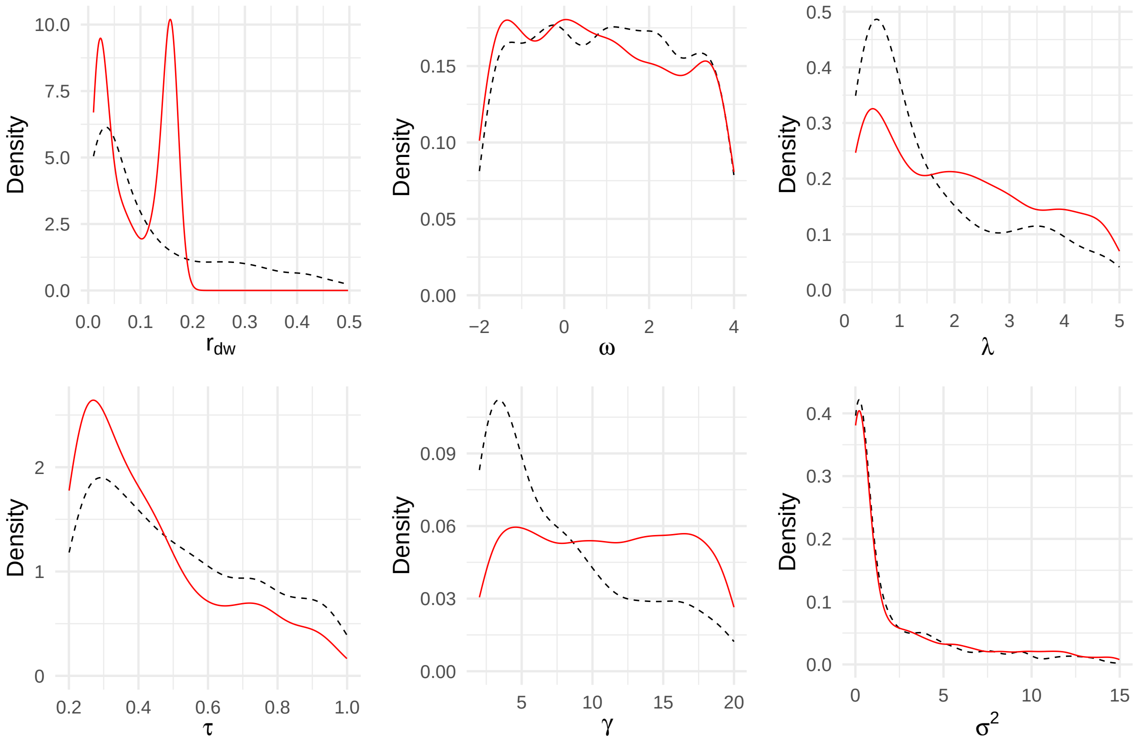


Figure S1: Distributions of parameters, before (in dotted black) and after (in red) retaining only parameters values leading to "realistic" epidemics. Dotted black distributions correspond to distributions given by Table S1. Red line distributions correspond to the distribution of parameters in $\theta_{\text{init}}$. We represent here the distribution of "realistic" epidemics from the four hypothetical dispersal models ($J_{\text{Exp}}$, $J_{\text{Gauss}}$, $J_{\text{ExpP}}$ and R.D.) for parameters $r_{\text{dw}}$, $\omega$, $\gamma$ and $\sigma^2$, for $J_{\text{ExpP}}$ for parameter $\tau$, and for $J_{\text{Exp}}$, $J_{\text{Gauss}}$ and $J_{\text{ExpP}}$ for parameter $\lambda$.

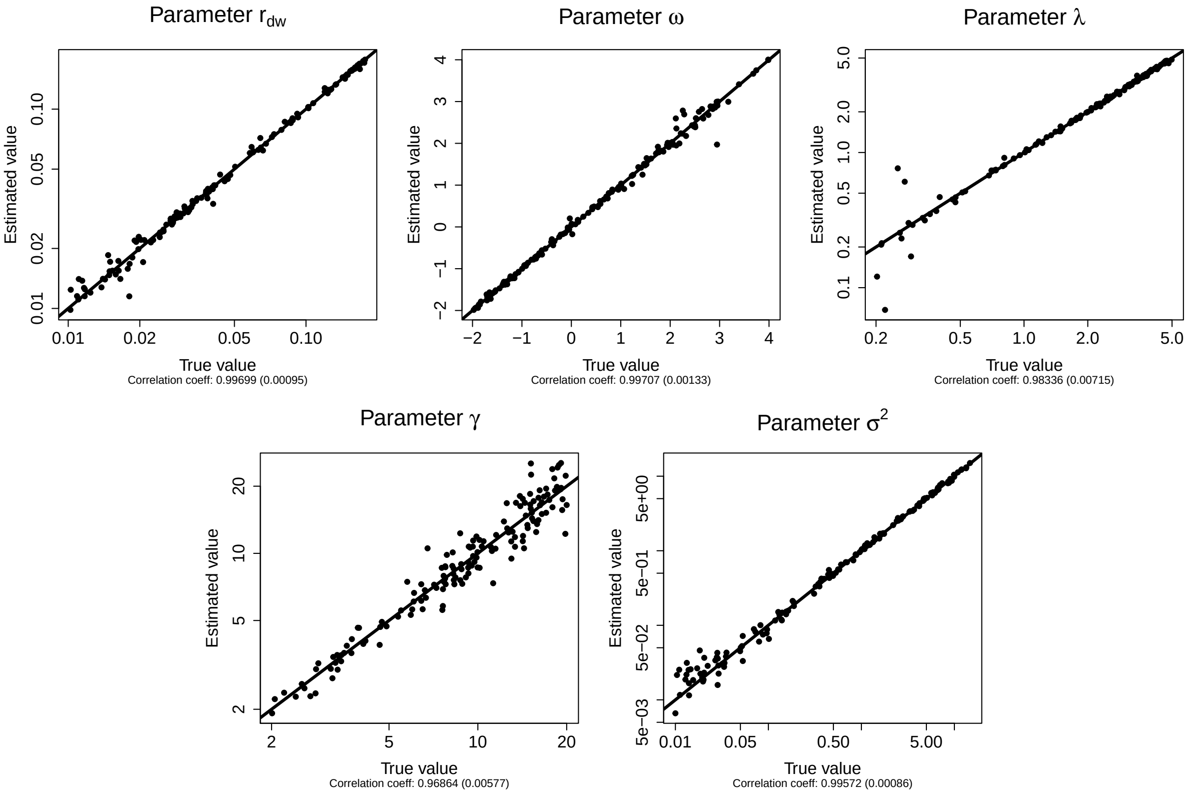


Figure S2: Practical parameter identifiability for the dispersal model $J_{\mathrm{Exp}}$. Each point represents the parameter estimation ('Estimated' value) depending on the real parameter ('True' value). Each graph regroups the results of 160 replicates. Straight lines correspond to the first bisector.

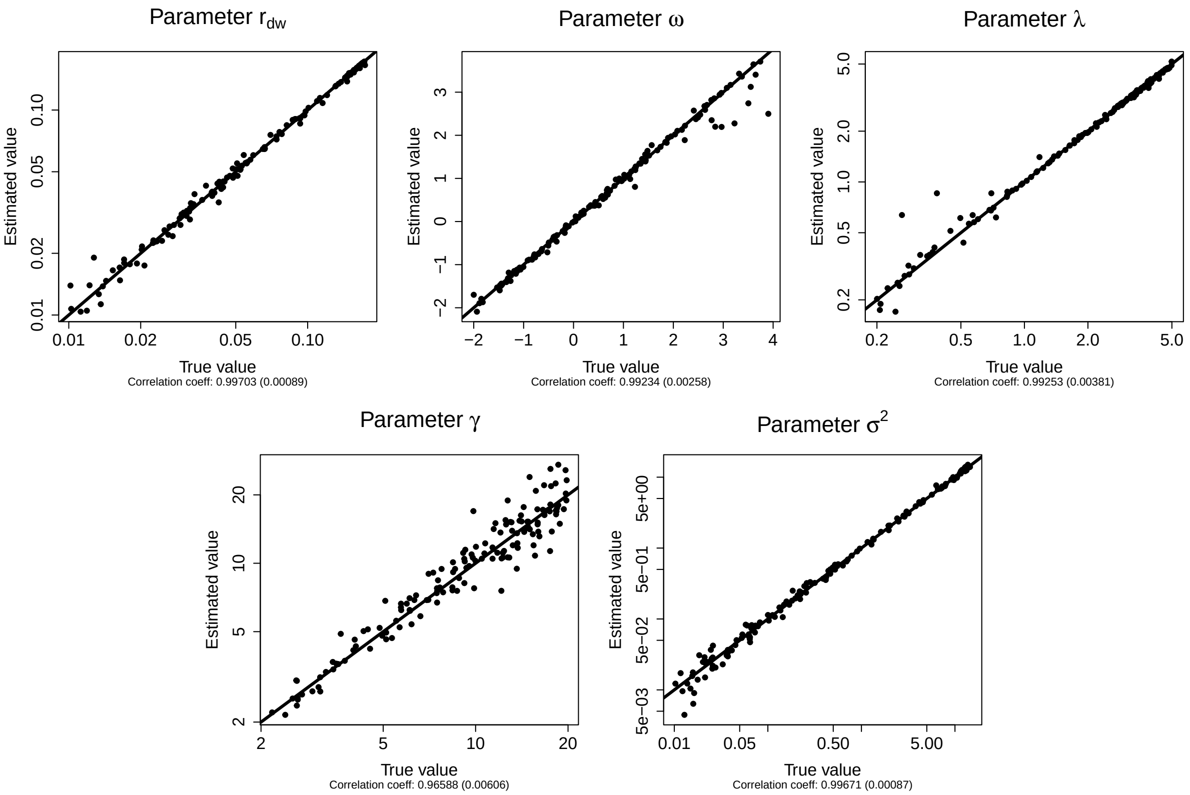


Figure S3: Practical parameter identifiability for the dispersal model $J_{\text{Gauss}}$. Each point represents the parameter estimation ('Estimated' value) depending on the real parameter ('True' value). Each graph regroups the results of 160 replicates. Straight lines correspond to the first bisector.

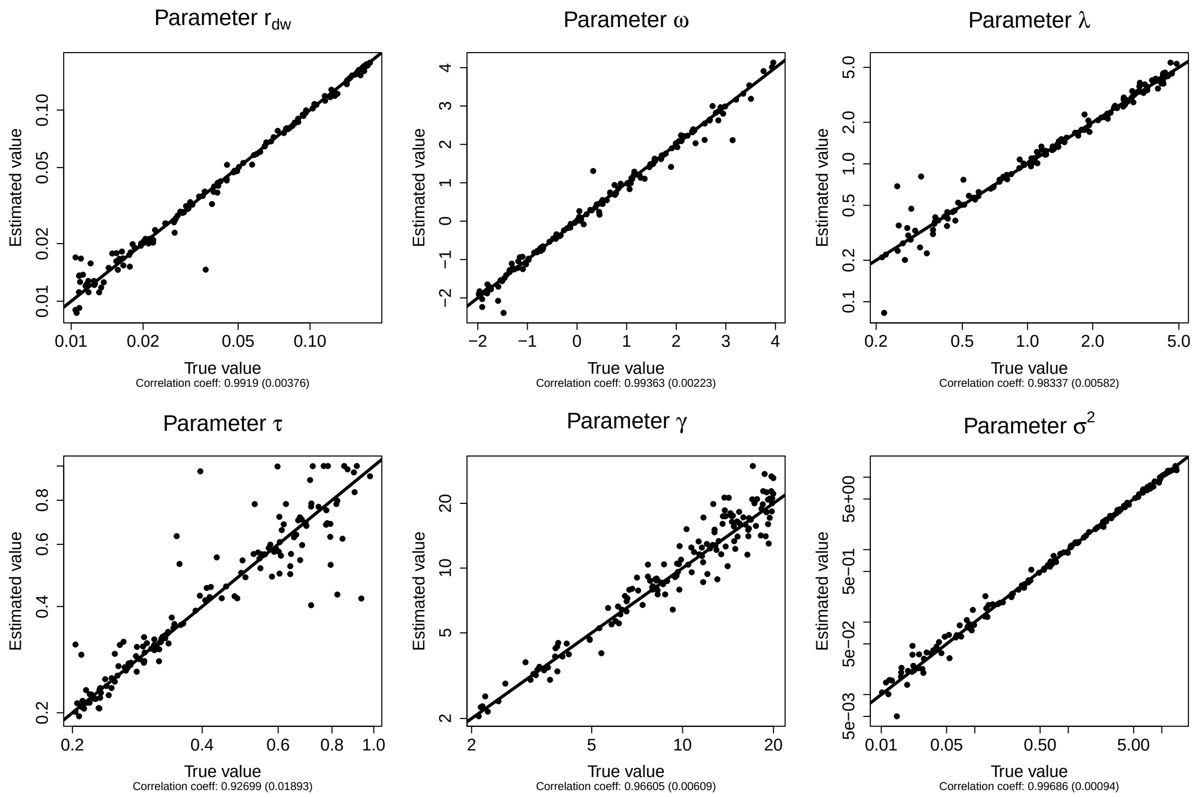


Figure S4: Practical parameter identifiability for the dispersal model $J_{\text{ExpP}}$. Each point represents the parameter estimation ('Estimated' value) depending on the real parameter ('True' value). Each graph regroups the results of 160 replicates. Straight lines correspond to the first bisector.

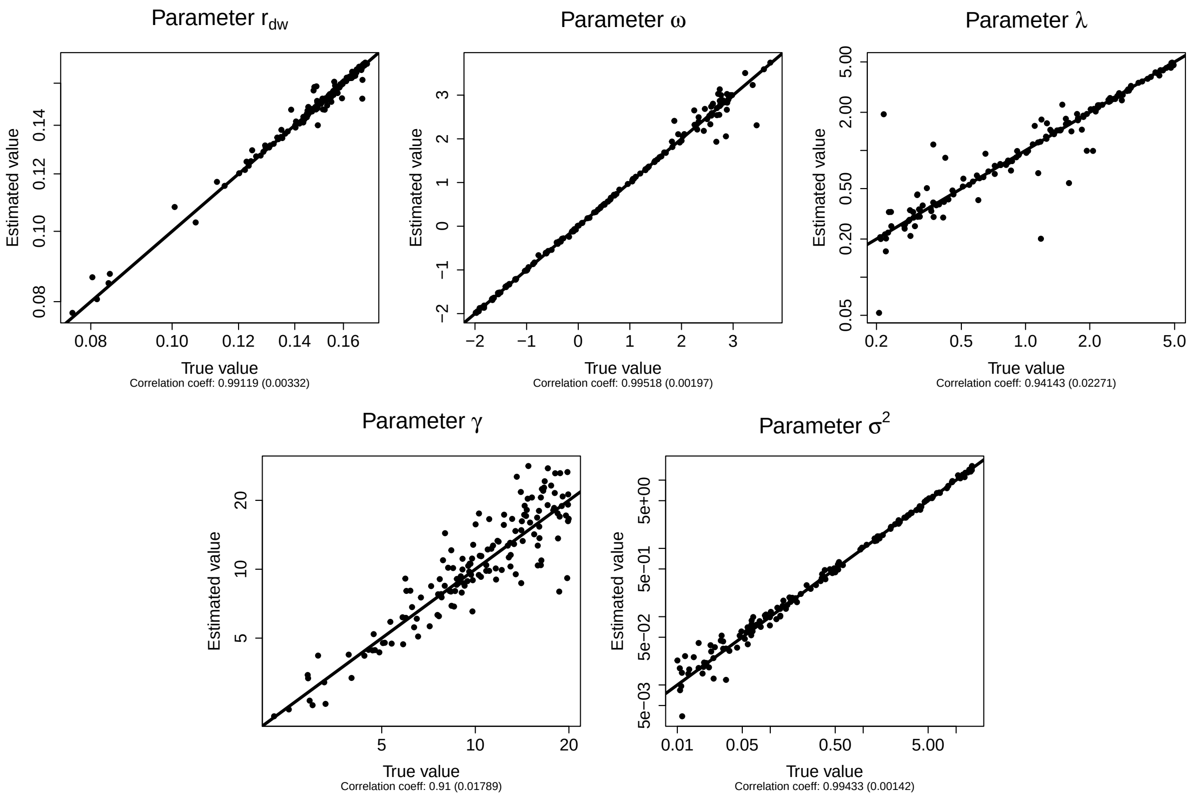


Figure S5: Practical parameter identifiability for the dispersal model R.D. Each point represents the parameter estimation (‘Estimated’ value) depending on the real parameter (‘True’ value). Each graph regroups the results of 160 replicates. Straight lines correspond to the first bisector.

#### S4.2 Model selection

Model practical identifiability was carried out in a similar way than parameter practical identifiability (Appendix S4.1), except that we fitted to each data set the true model (as previously) but also the three other models corresponding to the alternative hypotheses on the dispersal process. Models were compared using AIC (Akaike Information Criteria) to select the best data-supported model. AIC were assessed as $2k - 2ln(L)$ where $k$ is the number of parameters of the model considered and $L$ is the maximized value of the likelihood function. To gain more insights into the confidence level in model selection, we also calculated for each data set the difference between the AIC of the model selected and the AIC of the second-best model according to the two possible issues of the selection procedure: (i) when the model selection procedure was successful (*i.e.* the selected model was the true model) and (ii) when the model selection procedure was incorrect (*i.e.* the true model was not selected). The mean of these values were reported as $d\text{AIC}_{\text{true}}$ and $d\text{AIC}_{\text{wrong}}$ in Table 2. The steps were reiterated until the estimation of $n = 50$ realistic epidemics for each dispersal model.

#### S4.3 Parameter inference on the real data set

The model selection procedure was applied to the real data set by fitting four dispersal process hypotheses ($J_{\text{Exp}}$, $J_{\text{Gauss}}$, $J_{\text{ExpP}}$ and R.D.). The same optimisation routines described in Appendix S4.1 were performed from five initial parameter values selected as in Step 3.2 (Appendix S4.1). The selected model corresponds to hypothesis $J_{\text{ExpP}}$. For parameter estimations, we used the `mle2` function from the `R` package `bbmle`, with method NELDER-MEAD and optimizer NLMINB, to obtain maximum likelihood estimates of the vector of parameters $\hat{\theta}$ and of its matrix of variance-covariance $\hat{\Sigma}$. We used as initial parameter values the vector of parameters $\theta$ giving the lowest AIC value in the previous model selection procedure. Confidence intervals were derived from 1,000 random draws from the multivariate normal distribution with parameters $\hat{\theta}$ and $\hat{\Sigma}$. The 95% confidence intervals of each parameter is obtained using the quantiles 2.5% and 97.5% (Table 4).

#### S4.4 Model check

The model was checked by assessing the coverage rate of the data from the 95%-prediction intervals. The coverage rate was estimated as the proportion of observed data from the raw sampling within the prediction intervals (Figure 5).

Data from the raw sampling represent 97 counts $Y_{st}$ of infected trees at sites $s \in \{1,\dots,S\}$ (with $S = 12$ or $S = 45$ depending on the sampling date) and times $t \in \{1,\dots,6\}$. Let us recall that, as stated in Appendix S2, $Y_{st}$ follows a combination of Poisson and Beta-Binomial distributions whose parameters depend on the known mean value $(\lambda_m)_t$ and the unknown $u(t,x_s)$, $\gamma$ and $\sigma^2$, and that $u(t,x_s)$ is a deterministic function of dynamical parameters $r$, $\lambda$ and $\tau$.

Prediction intervals were calculated at each date and each site with a two-step procedure:

**Step 1** A confidence interval was obtained from 1000 random draws from the multivariate normal distribution with $\hat{\theta}$ and $\hat{\Sigma}$.

**Step 2** The mean proportions of infected trees were calculated at each date and site date from each random draw of parameters obtained from Step 1. A prediction interval was obtained from these parameters given the probabilities of infection, with 1,000 random draws in the observation laws.

Model checks were performed for each dispersal kernel model, and not only the selected model $J_{\text{ExpP}}$, to ensure that the coverage rates were higher with the selected model (Figure 5 for the selected dispersal model $J_{\text{ExpP}}$, and Figures S6 and S7 for dispersal models $J_{\text{Exp}}$ and $J_{\text{Gauss}}$, respectively). The model check was not performed for dispersal model R.D. because the estimated dispersal distance $\lambda_{\text{estim}}$ reached the upper limit of our numerical scheme $\lambda_{\text{up}} = 23$ and did not allow to calculate the confidence intervals.

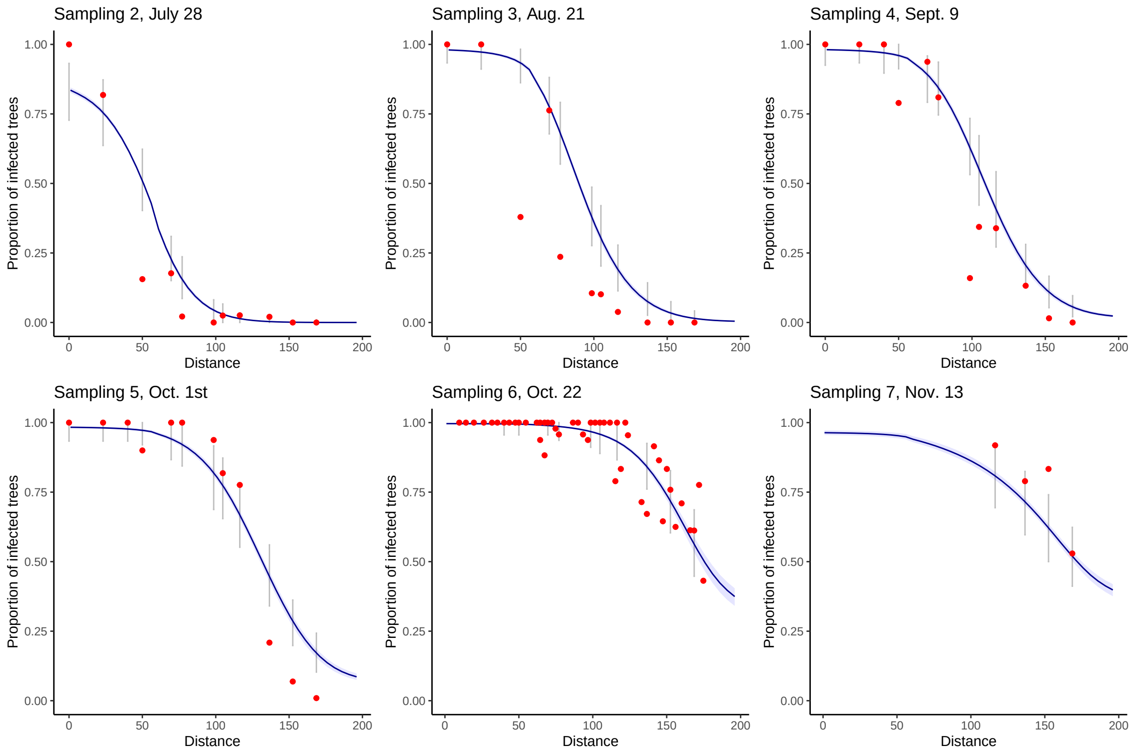


Figure S6: Model check under the dispersal model $J_{\text{Exp}}$: Coverage rates for the raw sampling. Each sampling date is represented on a separate graph. Sampling 1 is not represented because it corresponds to the initial condition of the epidemics for all simulations. Blue areas correspond to the pointwise 95% confidence envelopes for the proportion of infected trees, grey intervals correspond to the 95% prediction intervals at each site, *i.e.* taking into account the observation laws given the proportion of infected trees. Red points correspond to the observed data. Only four observations are available for sampling 7 because at this date (November 13) the leaves had already fallen from the trees located upstream the valley. The total coverage rate over all sampling dates is 0.68.

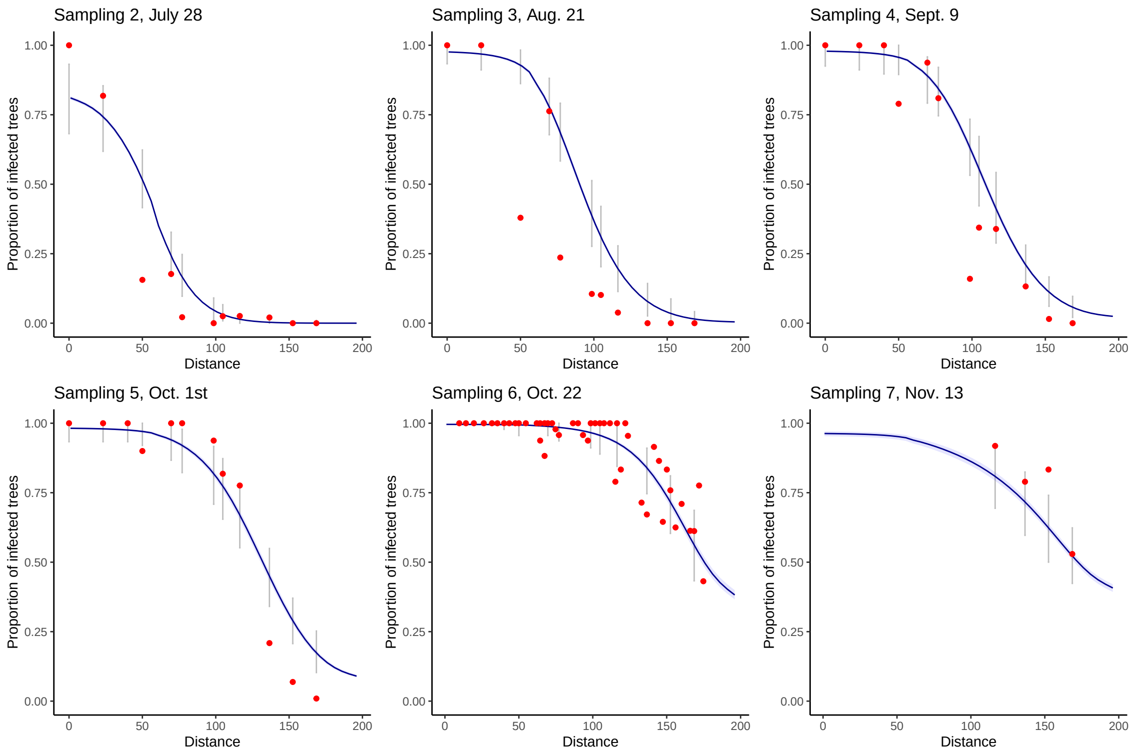


Figure S7: Model check under the dispersal model $J_{\text{Gauss}}$: Coverage rates for the raw sampling. Each sampling date is represented on a separate graph. Sampling 1 is not represented because it corresponds to the initial condition of the epidemics for all simulations. Blue areas correspond to the pointwise 95% confidence envelopes for the proportion of infected trees, grey intervals correspond to the 95% prediction intervals at each site, *i.e.* taking into account the observation laws given the proportion of infected trees. Red points correspond to the observed data. Only four observations are available for sampling 7 because at this date (November 13) the leaves had already fallen from the trees located upstream the valley. The total coverage rate over all sampling dates is 0.67.

#### S4.5 Sampling densification

As in Appendix S4.2, numerical simulations were run to disentangle the true dispersal process from alternative dispersal processes, with densification of time and site for the raw and the refined sampling. Simulations were run with 21 sampling dates instead of 6, which amounts to one sampling every week. The number of sampling sites was set to 45 for all sampling dates. The steps described in Appendix S4.1 and S4.2 were reiterated until the estimation of $n = 50$ realistic epidemics for each dispersal model.

Table S2: Efficiency of model selection for the densification of time samples (21 instead of 6) and the site sampled (45 instead of 12). The four first columns indicate the proportion of cases, among 50 replicates, where each tested model was selected using AIC, given that data sets were generated under a particular model (*i.e.* true model). Column $d\text{AIC}_{\text{true}}$ (*resp.* $d\text{AIC}_{\text{wrong}}$) indicates the mean difference between the AIC of the model selected when the model selected is the true one (*resp.* when the model selected is not the true model) and the second best model (*resp.* being the true model or not).

| | Selected Model | | | | | |
|---|---|---|---|---|---|---|
| True Model | $J_{\text{Exp}}$ | $J_{\text{Gauss}}$ | $J_{\text{ExpP}}$ | R.D. | $d\text{AIC}_{\text{true}}$ | $d\text{AIC}_{\text{wrong}}$ |
| $J_{\text{Exp}}$ | **0.64** | 0.18 | 0.12 | 0.06 | 5.60 | 2.02 |
| $J_{\text{Gauss}}$ | 0.14 | **0.8** | 0 | 0.06 | 9.93 | 1.51 |
| $J_{\text{ExpP}}$ | 0.1 | 0.02 | **0.86** | 0.02 | **2228.45** | 1.81 |
| R.D. | 0.12 | 0.16 | 0 | **0.72** | 32.97 | 1.04 |

### S5 Carrying capacity of poplar leaves

We measured the area of 10 wild poplar leaves (*Populus nigra*) and obtained a mean leaf area of $870\,mm^2$. We consider that poplar rust can not infect the leaf veins and edges, which represent approximately 15% of the leaf area. This leads to a net leaf area accessible to the pathogen of $740\,mm^2$. The size of a poplar rust lesion ranges from $0.2\,mm^2$ to $0.8\,mm^2$ (Maupetit et al., 2018). The lesions cannot fuse and are surrounded by living host tissue. We thus consider a lesion occupies a total area of $1\,mm^2$. This leads to a maximum of 740 lesions per leaf on average. To respect this order of magnitude, we consider in this analysis that the carrying capacity of a poplar leaf is 750 poplar rust lesions.
